# Positively selected modifications in the pore of TbAQP2 allow pentamidine to enter *Trypanosoma brucei*

**DOI:** 10.1101/2020.03.08.982751

**Authors:** Ali H. Alghamdi, Jane C. Munday, Gustavo D. Campagnaro, Dominik Gurvič, Fredrik Svensson, Chinyere E. Okpara, Arvind Kumar, Maria Esther Martin Abril, Patrik Milić, Laura Watson, Daniel Paape, Luca Settimo, Anna Dimitriou, Joanna Wielinska, Graeme Smart, Laura F. Anderson, Christopher M. Woodley, Siu Pui Ying Kelley, Hasan M.S. Ibrahim, Fabian Hulpia, Mohammed I. Al-Salabi, Anthonius A. Eze, Ibrahim A. Teka, Simon Gudin, Christophe Dardonville, Richard R Tidwell, Mark Carrington, Paul M. O’Neill, David W Boykin, Ulrich Zachariae, Harry P. De Koning

## Abstract

Mutations in the *Trypanosoma brucei* aquaporin AQP2 are associated with resistance to pentamidine and melarsoprol. We show that TbAQP2 but not TbAQP3 was positively selected for increased pore size from a common ancestor aquaporin. We demonstrate that TbAQP2’s unique architecture permits pentamidine permeation through its central pore and show how specific mutations in highly conserved motifs affect drug permeation. Introduction of key TbAQP2 amino acids into TbAQP3 renders the latter permeable to pentamidine. Molecular dynamics demonstrates that permeation by dicationic pentamidine is energetically favourable in TbAQP2, driven by the membrane potential, although aquaporins are normally strictly impermeable for ionic species. We also identify the structural determinants that make pentamidine a permeant but exclude most other diamidine drugs. Our results have wide-ranging implications for optimising antitrypanosomal drugs and averting cross-resistance. Moreover, these new insights in aquaporin permeation may allow the pharmacological exploitation of other members of this ubiquitous gene family.

## Introduction

The *Trypanosoma brucei*-group species are protozoan parasites that cause severe and fatal infections in humans (sleeping sickness) and animals (nagana, surra, dourine) (Giordani *et al,* 2016; Büscher *et al,* 2017). The treatment is dependent on the sub-species of trypanosome, on the host, and on the stage of the disease (Giordani *et al,* 2016; De Koning, 2020). Many anti-protozoal drugs are inherently cytotoxic but derive their selectivity from preferential uptake by the pathogen rather than the host cell (Munday *et al,* 2015a; De Koning, 2020). Conversely, loss of the specific drug transporters is a main cause for drug resistance (Barrett *et al,* 2011; Baker *et al,* 2013; Munday *et al,* 2015a; De Koning, 2020). This is the case for almost all clinically used trypanocides, including diamidines such as pentamidine and diminazene (Carter *et al,* 1995; De Koning, 2001a; De Koning *et al,* 2004; Bridges *et al,* 2007), melaminophenyl arsenicals such as melarsoprol and cymelarsan for cerebral stage human and animal trypanosomiasis, respectively (Carter & Fairlamb, 1993; Bridges *et al,* 2007), and the fluorinated amino acid analogue eflornithine for human cerebral trypanosomiasis (Vincent *et al,* 2010). The study of transporters is thus important for anti-protozoal drug discovery programmes as well as for the study of drug resistance (Lüscher *et al,* 2007; Munday *et al,* 2015a).

In *Trypanosoma brucei*, the phenomenon of melarsoprol-pentamidine cross-resistance (MPXR) was first described shortly after their introduction (Rollo & Williamson, 1951), and was linked to reduced uptake rather than shared intracellular target(s) (Frommel & Balber, 1987). The first transporter to be implicated in MPXR was the aminopurine transporter TbAT1/P2 (Carter & Fairlamb, 1993; Mäser *et al,* 1999; Munday *et al,* 2015b) but two additional transport entities, named High Affinity Pentamidine Transporter (HAPT1) and Low Affinity Pentamidine Transporter (LAPT1), have been described (De Koning, 2001a; De Koning & Jarvis, 2001; Bridges *et al,* 2007). HAPT1 was identified as Aquaglyceroporin 2 (TbAQP2) via an RNAi library screen, and found to be the main determinant of MPXR (Baker *et al,* 2012, 2013; Munday *et al,* 2014). The apparent permissibility for high molecular weight substrates by TbAQP2 was attributed to the highly unusual selectivity filter of TbAQP2, which lacks the canonical aromatic/arginine (ar/R) and full NPA/NPA motifs, resulting in a much wider pore (Baker *et al,* 2012; Munday *et al,* 2014, 2015a). Importantly, the introduction of TbAQP2 into *Leishmania* promastigotes greatly sensitised these cells to pentamidine and melarsen oxide (Munday *et al,* 2014). Moreover, in several MPXR laboratory strains of *T. brucei* the *AQP2* gene was either deleted or chimeric after cross-over with the adjacent *TbAQP3* gene, which, unlike AQP2, contains the full, classical ar/R and NPA/NPA selectivity filter motifs and is unable to transport either pentamidine or melaminophenyl arsenicals (Munday *et al,* 2014). Similar chimeric genes and deletions were subsequently isolated from sleeping sickness patients unresponsive to melarsoprol treatment (Graf *et al,* 2013; Pyana Pati *et al,* 2014) and failed to confer pentamidine sensitivity when expressed in a *tbaqp2-tbaqp3* null *T. brucei* cell line whereas wild-type TbAQP2 did (Munday *et al,* 2014; Graf *et al,* 2015).

The model of drug uptake through a uniquely permissive aquaglyceroporin (Munday *et al,* 2015a) was challenged by a study arguing that instead of traversing the TbAQP2 pore, pentamidine merely binds to an aspartate residue (Asp265) near the extracellular end of the pore, above the selectivity filter, followed by endocytosis (Song *et al,* 2016). This alternative, ‘porin-receptor’ hypothesis deserves careful consideration given that (i) it is an exceptional assertion that drug-like molecules with molecular weights grossly exceeding those of the natural substrates, can be taken up by an aquaglyceroporin and (ii) the fact that bloodstream form trypanosomes do have, in fact, a remarkably high rate of endocytosis (Field & Carrington, 2009; Zoltner *et al,* 2016). The question is also important because aquaporins are found in almost all cell types and the mechanism by which they convey therapeutic agents and/or toxins into cells is of high pharmacological and toxicological interest. While TbAQP2 is the first aquaporin described with the potential to transport drug-like molecules, this ability might not be unique, and the mechanism by which the transport occurs should be carefully investigated.

We therefore conducted a mutational analysis was undertaken, swapping TbAQP2 and TbAQP3 selectivity filter residues and altering pore width at its cytoplasmic end. This was complemented with a thorough structure-activity relationship study of the interactions between pentamidine and TbAQP2, using numerous chemical analogues for which inhibition constants were determined and interaction energy calculated. The pentamidine-TbAQP2 interactions were further modelled by running a molecular dynamics simulation on a protein-ligand complex, and In addition, we investigated a potential correlation between the *T. brucei* endocytosis rate and the rate of pentamidine uptake. Our results unequivocally show that pentamidine permeates directly through the central pore of TbAQP2 and that uptake is dependent on the microbial membrane potential. Having identified the essential characteristics that allow the transport of large, flexible molecules through TbAQP2, this should now allow the evaluation of aquaporins in other species for similar adaptations.

## Results

### 12. Investigation of the structural determinants of AQP2 for pentamidine transport

#### 1.1. Positive selection for pore size

In *T. brucei*, the AQP2 and AQP3 genes are arranged as a tandem pair on chromosome 10 and have 74% amino acid identity. Whereas TbAQP2 clearly mediates pentamidine uptake, TbAQP3 does not (Baker *et al,* 2012; Munday *et al,* 2014), nor do various chimeric AQP2/3 rearrangements that give rise to pentamidine resistance (Munday *et al,* 2014; Graf *et al,* 2015). To investigate the origin of the AQP2 gene, a phylogenetic analysis of AQPs in African trypanosomes was performed. The number of aquaporin genes varies: there is a single aquaporin in *T. vivax* and *T. congolense,* two in *T. suis* and three in *T. brucei* and its derivatives (Supplemental Fig. 1A). The most probable tree (Supplemental Fig. 1B) is consistent with the evolutionary history of the four species (Hutchinson & Gibson, 2015) and indicates AQP1 as the ancestral AQP present in all trypanosome species. A duplication occurred in the common ancestor of *T. suis* and *T. brucei* after divergence from *T. congolense* and a further duplication, to form AQP2 and AQP3, in the ancestor of *T. brucei* after divergence from *T. suis*. Multiple alignment (Supplemental Fig. 1A) shows that the classical NPA/NPA and ar/R AQP selectivity filter elements are present in all AQPs except *T. brucei* AQP2. The divergence of *T. brucei* AQP2 and 3 was investigated by calculating the non-synonymous/synonymous codon ratio (dN/dS) for different AQPs (Supplemental Fig. 1C). For *T. brucei* aligned with *T. suis* AQP1, dN/dS is 0.21 and for AQP3, dN/dS is 0.30 indicating purifying selection. However, comparing *T. brucei* AQP2 with *T. brucei* AQP3, dN/dS is 2.0 indicating strong selection pressure for divergence on AQP2 towards an aquaporin with increased pore size. In order to verify any role of amino acids along the TbAQP2 pore in facilitating pentamidine sensitivity and/or uptake, we performed a mutational analysis.

#### 1.2 Introduction of AQP3 residues into the AQP2 selectivity filter

One highly conserved motif of aquaporins, believed to be essential for permeant selectivity, is NPA/NPA which is present in TbAQP3 but not in TbAQP2, where, uniquely, it is N**S^131^**A/NP**S^263^** instead. We therefore constructed a TbAQP2 variant with the classical NPA/NPA motif (TbAQP2^S131P/S263A^) and expressed it in the *aqp2/aqp3* null cell line (Baker *et al,* 2012; Munday *et al,* 2014). In this cell line, uptake of 30 nM [^3^H]-pentamidine was reduced to 4.40 ± 0.71% (*n*=4) of the rate in the control line expressing TbAQP2WT (P<0.05, Student’s unpaired t-test), as well as significantly different from the rate measured in parallel in the *tbaqp2/tbaqp3* null cells (P<0.01) (Fig. 1A). The remaining pentamidine uptake was sufficient to strongly sensitise the TbAQP2^S131P/S263A^ cells to pentamidine in a standard protocol of 48 h incubation with the drug followed by a further 24 h in the presence of the resazurin indicator dye (P<0.0001 vs *tbaqp2/tbaqp3* null) but the EC_50_ was still significantly higher than the TbAQP2WT control (P<0.05) (Fig. 1B). A similar effect was observed for the melaminophenyl arsenical drug cymelarsan, but there was no change in sensitivity to diminazene or the control drug phenylarsine oxide (PAO), which is believed to diffuse directly across the membrane (Fairlamb *et al,* 1992) (Fig. 1B).

**Fig. 1.**
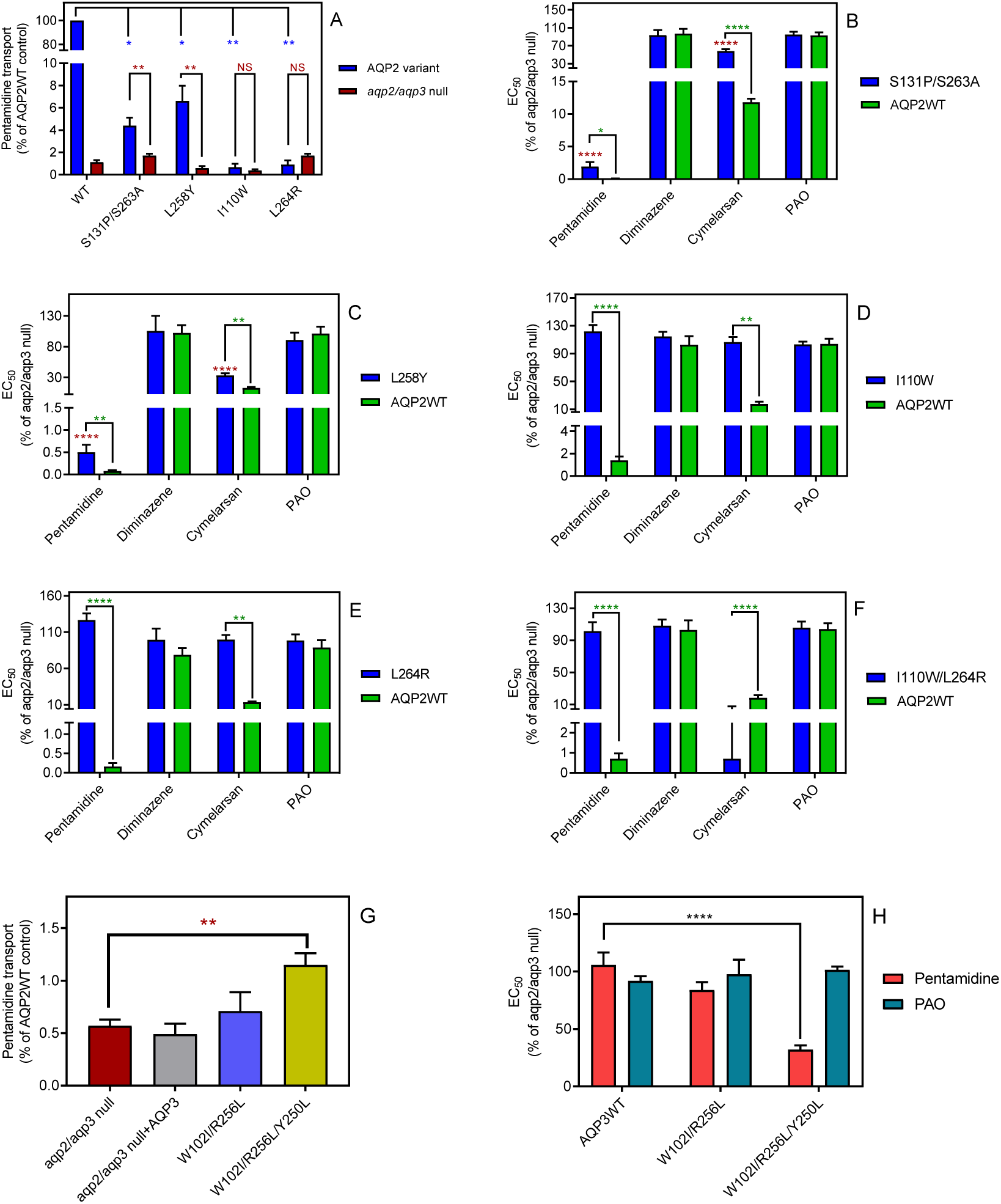
The selectivity filter differences between TbAQP2 and TbAQP3 are largely responsible for their differences in pentamidine sensitivity and transport rates. (A) Transport of 30 nM [^3^H]-pentamidine by *tbaqp2/aqp3* null cells expressing TbAQP2-WT or one of the TbAQP2 mutants as indicated (blue bars). The corresponding brown bars are pentamidine transport in the control *tbaqp2/aqp3* null cells assessed in parallel in each experiment. Transport was determined in the presence of 1 mM adenosine to block the TbAT1/P2 transporter. Bars represent the average and SEM of at least three independent experiments, each performed in triplicate. Blue stars: statistical significance comparison, by two-tailed unpaired Student’s tests, between the cells expressing TbAQP2WT and mutants; red stars: statistical comparison between the AQP2-expressing cells and control cells; NS, not significant. (B-F) EC_50_ values indicated test drugs, expressed as a percentage of the resistant control (*tbaqp2/tbaqp3* null), against cell lines either expressing the indicated TbAQP2 mutant or TbAQP2WT (sensitive control). Red stars and green stars: comparison with *tbaqp2/aqp3* null or TbAQP2WT-expressing cells, respectively, which were always assessed in parallel in each experiment. (G) Transport of 30 nM [^3^H]-pentamidine by *tbaqp2/aqp3* null cells expressing TbAQP3 or an AQP3 mutant as indicated. (H) EC_50_ values of the indicated drugs against *tbaqp2/aqp3* null cells expressing either TbAQP3 or a mutant thereof, expressed as percentage of *tbaqp2/aqp3* null. All experiments are the average and SEM of at least 3 independent experiments. *, P<0.05; **, P<0.01; ***, P<0.001, ****, P<0.0001 by unpaired Student’s t-test, two-tailed.

The mutant L258Y, which has the AQP3 Tyr-250 half of the highly conserved aromatic/arginine (ar/R) motif, responsible for pore restriction and proton exclusion (Wu *et al,* 2009), introduced into the TbAQP2 pore, yielded a drug transport phenotype similar to TbAQP2^S131P/S263A^. The [^3^H]-pentamidine transport rate was reduced to 6.6 ± 1.4% of TbAQP2WT (P<0.05) but remained above the rate in the *tbaqp2/tbaqp3* null cells (P<0.01) (Fig. 1A). Pentamidine and cymelarsan EC_50_ values were also significantly different from both the TbAQP2WT and the *tbaqp2/tbaqp3* null controls (Fig. 1C).

The ar/R motif is part of the larger selectivity filter, usually WGYR, present in both TbAQP1 and TbAQP3 but uniquely consisting of I^110^VL^258^L^264^ in TbAQP2 (Baker *et al,* 2013), all non-polar, open chained residues. Cell lines expressing mutations AQP2^I110W^ and AQP2^L264R^, either alone or in combination, displayed pentamidine transport rates, and pentamidine and cymelarsan EC_50_ values that were not significantly different from the *tbaqp2/tbaqp3* null controls but highly significantly different from the TbAQP2WT drug-sensitive controls, showing that their capacity for pentamidine and cymelarsan uptake had been reduced to zero (Fig. 1A, D-F).

We conclude that the unique TbAQP2 replacement of the NPA/NPA motif and all of the WGYR selectivity filter mutations are necessary for the observed pentamidine and melaminophenyl arsenical sensitivity observed in cells expressing wild-type TbAQP2.

#### 1.3 Introduction of TbAQP2 selectivity filter residues into the AQP3 pore enables pentamidine transport

An interesting question was whether the introduction of (some of) the critical TbAQP2 residues in TbAQP3 might give the latter the capacity to take up pentamidine. We therefore constructed TbAQP3^W102I/R256L^ and TbAQP3^W102I/R256L/Y250L^ and tested whether *tbaqp2/tbaqp3* null cells transfected with these mutant aquaporins were able to take up 25 nM [^3^H]-pentamidine in the presence of 1 mM adenosine (which blocks uptake via TbAT1/P2). Pentamidine uptake in the tested cell lines was very low compared to the same cells expressing TbAQP2WT (Fig. 1G). However, by measuring [^3^H]-pentamidine uptake over 30 min it was possible to reliably and reproducibly measure radiolabel accumulation in each cell line. This showed that while uptake in TbAQP3^W102I/R256L^ only trended slightly upwards (P>0.05), the mutant AQP3 with all three AQP2 WGYR residues (W102I, R256L and Y250L) accumulated significantly more [^3^H]-pentamidine than the *tbaqp2/tbaqp3* null cells (P<0.01) or the null cells expressing TbAQP3WT (P=0.011). This is further corroborated by comparing the pentamidine sensitivity profile of these cell lines: only TbAQP3^W102I/R256L/Y250L^ conveyed significant sensitisation to *tbaqp2/tbaqp3* null cells (P<0.0001; Fig. 1H). Thus, TbAQP3 is converted into a pentamidine transporter by the insertion of the AQP2 WGYR residues, although this does not convey as high a rate of pentamidine uptake as TbAQP2.

#### 1.4 Mutations of amino acids modelled to potentially bind pentamidine or melarsoprol dramatically reduce pentamidine transport

Our previous attempts at modelling the binding of pentamidine and melarsoprol into the pore of TbAQP2 tentatively identified several residues that could be involved in this process (Munday *et al,* 2015a), from which we selected two residues, Ile190 and Trp192, at the extracellular end of the channel (position shown in Fig. 6), to swap with the corresponding residues of TbAQP3, creating TbAQP2^I190T^ and TbAQP2^W192G^. Both residues were predicted to interact with the substrate(s) via main-chain carbonyl oxygen atoms, but the side chains could nonetheless affect the interactions.

TbAQP2^I190T^ displayed dramatically reduced [^3^H]-pentamidine uptake, at 2.7 ± 0.7% (P<0.01, *n*=4) of the TbAQP2WT control, although significantly higher than the rate of the *tbaqp2/tbaqp3* null negative control (P<0.05) (Fig. 2A). The reduced rate was the result of a reduced V_max_ of the high affinity [^3^H]-pentamidine uptake, rather than a change in Km; the LAPT1 V_max_ and K_m_ were unchanged in cells expressing TbAQP2^I190T^ or TbAQP2WT (Supplemental Fig. S2). TbAQP2^I190T^ still conferred some increased pentamidine sensitivity in the standard resazurin test (P<0.0001), although highly significantly less sensitizing than TbAQP2WT (P<0.001); an intermediate sensitivity was also observed for cymelarsan (Fig. 2B). Substitution W192G also produced intermediate sensitivity to both drugs (Fig. 2C) but the double substitution TbAQP2^I190T/W192G^ displayed no significant pentamidine uptake above *tbaqp2/tbaqp3* null (Fig. 3A) and did not sensitise to pentamidine or cymelarsan (Fig. 2D).

**Fig. 2.**
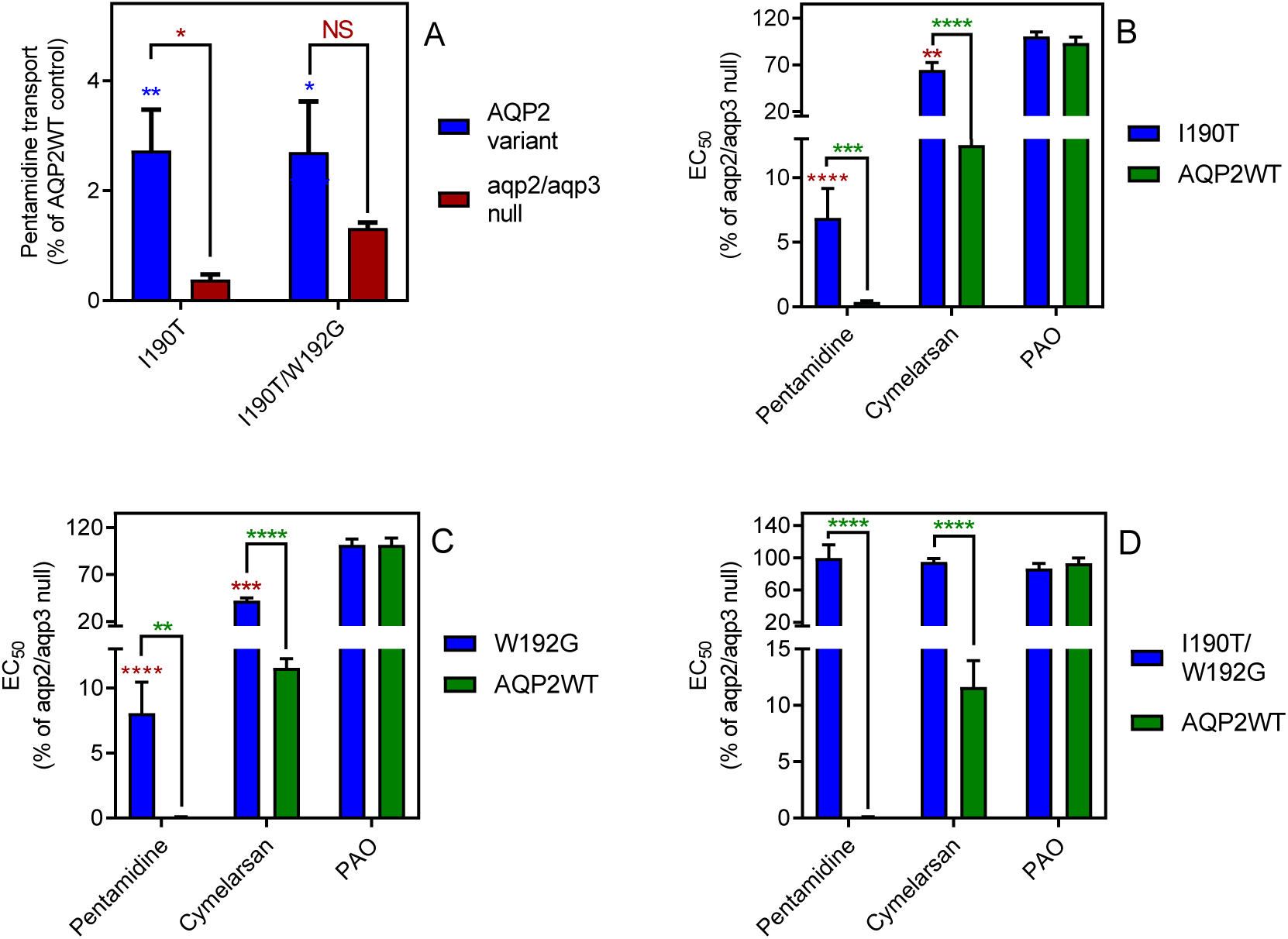
Mutational analysis of TbAQP2 residues I190 and W192. (A) Transport of 30 nM [^3^H]-pentamidine by *tbaqp2/tbaqp3* null cells or TbAQP2 variants expressed therein. Transport was expressed as a percentage of the rate of the AQP2WT control, performed in parallel. Blue stars are comparison with TbAQP2WT, red stars, comparison with the *tbaqp2/tbaqp3* null control. NS, not significant. (B) EC_50_ values for the indicated drugs against *tbaqp2/tbaqp3* null cells, and against TbAQP2WT and TbAQP2^I190T^ expressed therein; values were expressed as % of the *tbaqp2/tbaqp3* null (resistant) control. Red stars, comparison with the resistant control; green stars, comparison with the internal sensitive control (TbAQP2WT). The assays for all three strains and all three drugs were done simultaneously on at least 3 different occasions. (C) As B but for TbAQP2^W192G^. (D) As B but for TbAQP2^I190T/W192G^. *, P<0.05; **, P<0.01; ***, P<0.001, ****, P<0.0001 by unpaired Student’s t-test.

**Fig. 3.**
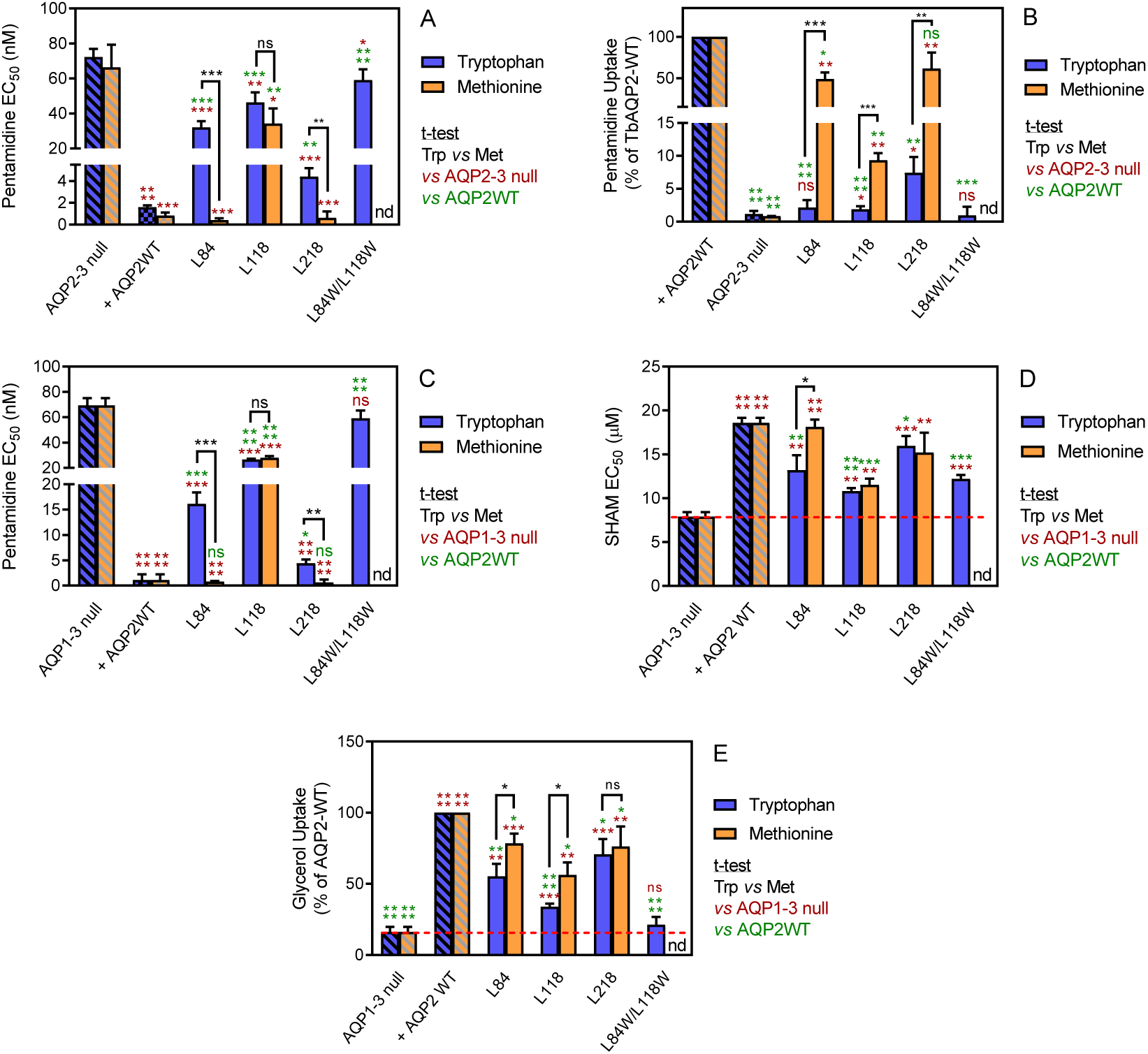
Analysis of TbAQP2 variants with a leucine-to-tryptophan or leucine-to-methionine substitution near the cytoplasmic end of the pore. (A) Pentamidine EC_50_ values (nM) for mutant and WT TbAQP2 expressed in *tbaqp2/tbaqp3* cells (aqp2-3 null). The mutants are either a Trp (dark blue bars) or Met (orange bars) substitution at the indicated positions. The resistant control (aqp2-3 null) and sensitive control (AQP2WT) for the separate datasets (Trp or Met) are indicated as hatched bars in the same colours. (B) As (A) but showing transport of 30 nM [^3^H]-pentamidine by the same cell lines, expressed as percentage of the transport rate in the TbAQP2 control cells. (C) Pentamidine EC_50_ values for the same mutants as in (A) but expressed in the *tbaqp1-2-3* null cells, performed in parallel with the determination of EC_50_ values for SHAM, shown in (D). As all cell lines were done simultaneously, the resistant and sensitive strain control values are identical for the Trp and Met mutants in this series. All bars represent the average and SEM of at least three independent replicates. *, P<0.05; **, P<0.01; ***, P<0.001, ****, P<0.0001 by unpaired Student’s t-test; ns, not significant; nd, not determined.

#### 1.5. The effect of large amino acids at the cytoplasmic end of the pore

To test whether restrictions at the cytoplasmic end of TbAQP2 would impact on pentamidine transport, we selected three leucine residues and exchanged each with tryptophan, creating L84W, L118W and L218W (positions indicated in Fig. 6). Expressing each of the L-to-W mutants in *tbaqp2/tbaqp3* null cells revealed significantly reduced pentamidine sensitivity compared to the same cells expressing TbAQP2WT (Fig. 3A), while also exhibiting dramatically reduced rates of [^3^H]-pentamidine transport (Fig. 3B). This effect was additive, with TbAQP2^L84W/L118W^ not significantly sensitising for pentamidine and displaying no detectable increase in [^3^H]-pentamidine transport relative to *tbaqp2/tbaqp3* null cells (Fig. 3A,B). None of these L-to-W mutants sensitised the cells to cymelarsan, diminazene or PAO (Supplemental Fig. S3). When the same leucine residues were replaced with methionine instead of tryptophan, variants L84M and L218M were not or barely different from TbAQP2WT with respect to pentamidine sensitisation (Fig. 3A) or transport (but highly significantly different from their respective tryptophan variants). For position 118 the Met replacement had similar effects as the Trp variant had, albeit with a significantly higher rate of pentamidine transport (1.88 ± 0.20 (*n*=6) *versus* 9.38 ± 0.63% (*n*=3) of TbAQP2WT, P<0.001; Fig. 3A,B). The L84M and L218M mutants also sensitised to cymelarsan (P<0.01) and, surprisingly, the L218 W and M mutants also sensitised slightly to diminazene (∼2-fold, P<0.05) (Supplemental Fig. S3).

These results strongly argue that the introduction of large amino acids at the cytosolic end significantly blocks the transport of pentamidine, whereas the change to Leu◊Met mutants were more permissive for pentamidine, but not cymelarsan. In order to check whether these variants were still functional aquaglyceroporins, we used the observation of Jeacock *et al,* (2017) that *T. brucei* cells lacking all three AQPs are sensitised to the Trypanosome Alternative Oxidase inhibitor SHAM, as a result of cellular glycerol accumulation. By this measure, all of the position 84, 118 and 218 Trp and Met mutants were able to transport glycerol, as each displayed SHAM EC_50_ values significantly different from the *tbaqp1-2-3* null cells (Fig. 3C,D); several variants displayed an intermediate SHAM EC_50_, being also significantly different from TbAQP2WT, indicating some attenuation of glycerol efflux capacity for those mutants. Indeed, uptake of [^3^H]-glycerol closely mirrored the SHAM observations (Fig. 3E).

#### 1.6. Overall correlation between [^3^H]-pentamidine transport rate and pentamidine EC_50_

The results presented in Figures 2-4 consistently show that even TbAQP2 mutants that display a large reduction in [^3^H]-pentamidine uptake rate results can show intermediate pentamidine sensitivity phenotypes (EC_50_s), due to the nature of the standard drug sensitivity test employed, which involves a 48-h incubation with the drug prior to a further 24-h incubation with resazurin: even a much-reduced transport rate will be sufficient to accumulate significant amounts of intracellular pentamidine over 3 days. A plot of [^3^H]-pentamidine transport rates versus pentamidine EC_50_, using the data for all 19 TbAQP2 and TbAQP3 mutants for which the transport rates were determined, shows that relatively small changes in EC_50_ occur, even with up to approximately 95% reduction in transport rates; at >95% reduction large EC_50_ increases become apparent (Supplemental Fig. S4).

**Fig. 4.**
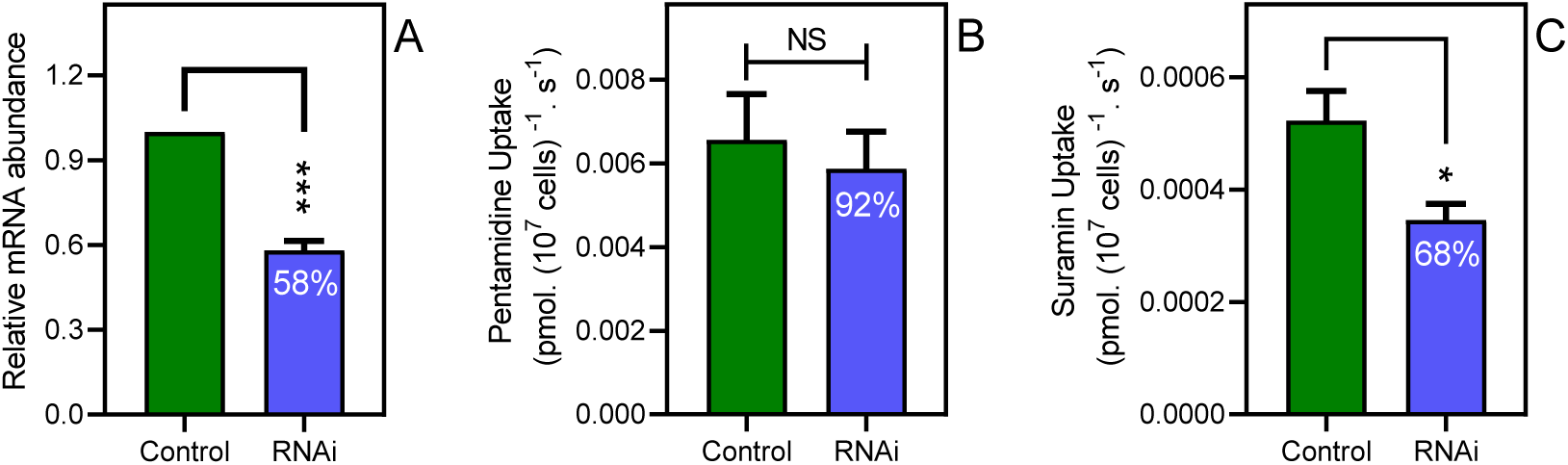
Disabling endocytosis does not reduce uptake of pentamidine. (A) qRT-PCR of CRK12, normalised to housekeeping gene GPI-8 (n=3). (B) Transport of 0.025 µM [^3^H]-Pentamidine measured in control (non-induced) and CRK12 cell after exactly 12 h of tetracycline induction; incubation time with label was 30 s. Bar is average and SEM of 5 independent determinations, each performed in triplicate. NS, not significant by unpaired Student’s t-test. (C) As frame B but uptake of 0.25 µM [^3^H]-suramin over 15 min; average and SEM of 5 independent determinations, each in quadruplicate. **, P=0.0027 by Student’s unpaired, two-tailed t-test.

### 2. Partially blocking endocytosis does not alter the rate of pentamidine transport

The knockdown of the CRK12 kinase in *T. brucei* causes a highly reproducible defect in endocytosis that affects an estimated one third of cells 12 h after RNAi induction and is ultimately lethal (Monnerat *et al,* 2013). We utilized this system to investigate whether a link between endocytosis and pentamidine transport exists. At 12 h of CRK12 RNAi induction with tetracycline, CRK12 mRNA levels were reduced by 42% (P<0.001) relative to uninduced controls as determined by qRT-PCR (Fig. 4A). Samples from the culture taken at this time point showed an increased abundance of cells with swelling characteristic of endocytosis defects, although this was hard to quantify as a minority of cells were affected, and to various degrees, as the 12 h time point was deliberately taken as an early point that would not yet affect cell viability (Supplemental Fig. S5) or cause excessive cellular pathology. We thus performed parallel uptake experiments with [^3^H]-pentamidine and [^3^H]-suramin, with suramin acting as positive control as it is known to enter *T. brucei* bloodstream forms through endocytosis after binding to surface protein ISG75 (Zoltner *et al,* 2016). After 12 h of CRK12 RNAi induction, pentamidine uptake was not significantly less than in the *T. brucei* 2T1 parental cells, whereas uptake of [^3^H]-suramin was (P=0.019, *n*=5; Fig. 4B,C).

### 3. The protonmotive force drives AQP2-mediated pentamidine uptake in bloodstream forms of T. brucei

It has been reported that knock-down of the HA1–3 plasma membrane proton pumps of *T. brucei* (which are essential for maintaining the plasma membrane potential), confers pentamidine resistance (Alsford *et al,* 2012; Baker *et al,* 2013). Interestingly, this locus only conferred resistance to (dicationic) pentamidine, not to the (neutral) melaminophenyl arsenicals, unlike knockdown of the TbAQP2/TbAQP3 locus (Alsford *et al,* 2012). We have previously reported that the HAPT-mediated pentamidine uptake in *T. brucei* procyclics correlates strongly with the proton-motive force (PMF) (De Koning, 2001a). However, it is not clear whether this dependency indicates that pentamidine uptake is mediated by a proton symporter, as known for many *T. brucei* nutrient transporters (De Koning & Jarvis, 1997a,b, 1998; De Koning *et al,* 1998), or reflects the energetics of uptake of cationic pentamidine being driven by the strong inside-negative membrane potential V_m_. The absence of an effect of HA1–3 knockdown on sensitivity to the neutral melaminophenyl arsenicals strongly argues against a mechanism of proton symport for HAPT1/AQP2 but a (partial) dependency of HAPT1/AQP2-mediated uptake of dicationic pentamidine on PMF or V_m_ would be expected if the substrate traverses the channel, as opposed to binding a single Asp residue on the extracellular side of the protein, as suggested in the endocytosis model (Song *et al,* 2016). Here we show that the same ionophores that inhibit HAPT1-mediated pentamidine transport in procyclic cells, and inhibit hypoxanthine uptake in both bloodstream form (BSF) (De Koning & Jarvis, 1997b) and procyclic (De Koning & Jarvis, 1997a) *T. brucei*, also dose-dependently inhibit [^3^H]-pentamidine uptake in BSF (Fig. 5A). This confirms that pentamidine needs the membrane potential for rapid uptake, as predicted by the dependence on the HA1–3 proton pumps. Using [^3^H]-suramin as an endocytosed substrate (Zoltner *et al,* 2016), we found that 20 µM CCCP also inhibits endocytosis in *T. brucei*, by 32.6% (P=0.029; pre-incubation 3 min, plus suramin accumulation over 10 minutes) (Fig. 5B). While that means that the ionophore experiments do not perfectly discriminate between endocytosis and trans-channel transport for di-cationic pentamidine, they do for neutral melaminophenyl arsenicals: the non-dependence of these neutral TbAQP2 substrates on the proton gradient (Alsford *et al,* 2012) indicates that, unlike suramin, they are not endocytosed.

**Fig. 5.**
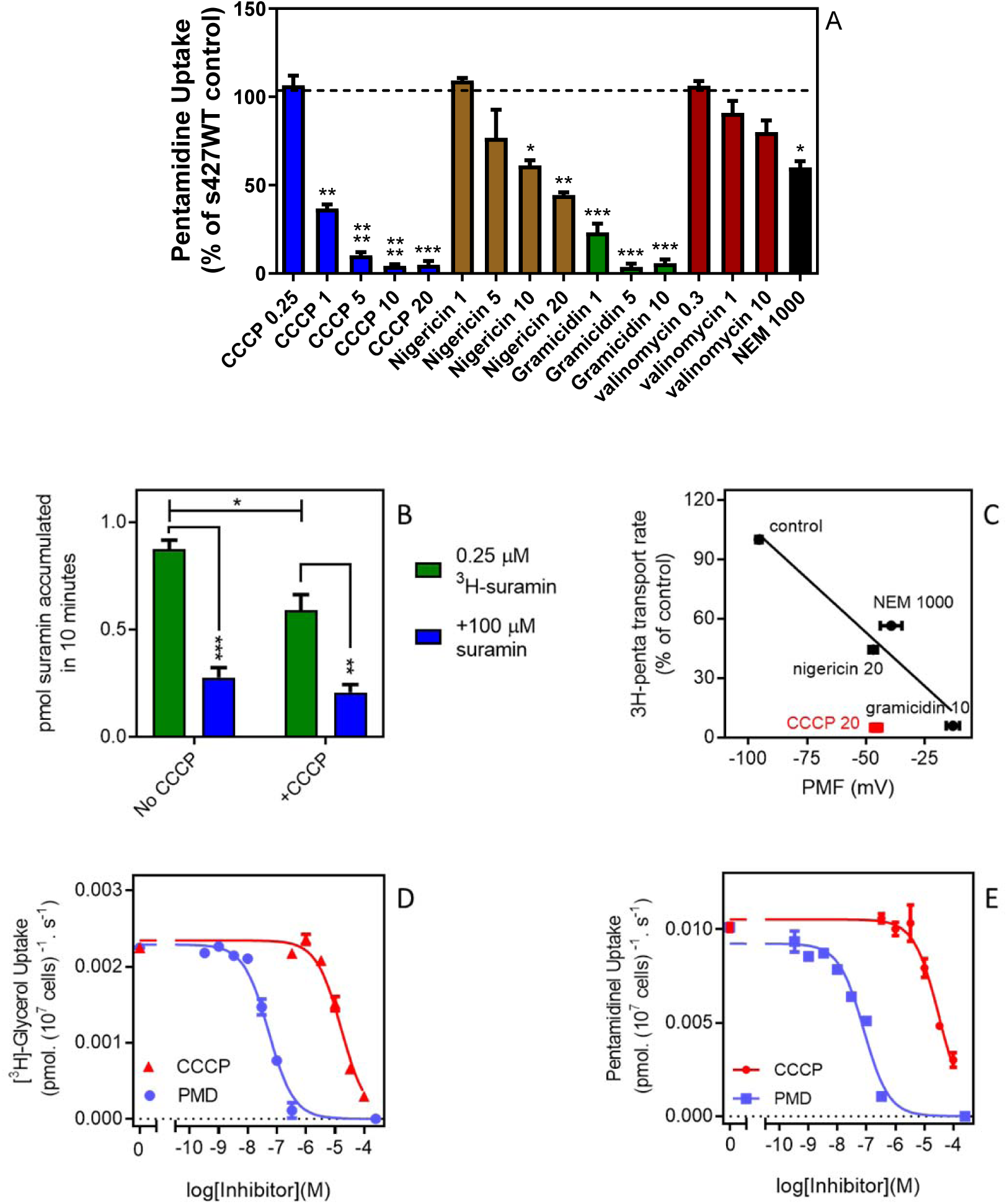
High affinity pentamidine uptake in *T. b. brucei* is sensitive to ionophores. (A) Uptake of 25 nM [^3^H]-pentamidine in s427WT bloodstream forms was measured in the presence of 1 mM adenosine to block the P2 transporter, and in the further presence of various ionophores at the indicated concentrations in µM. Incubation with radiolabel was 5 min after a 3 min pre-incubation with ionophore. Accumulation of radiolabel was expressed as a percentage of the control, being a parallel incubation in the absence of any ionophore. Bars represent the average of 3 – 5 independent determinations (each performed in quadruplicate) and SEM. (B) Uptake of 0.25 µM [^3^H]-suramin by *T. b. brucei* s427WT cells over 10 minutes. Cells were incubated in parallel, with or without the presence of 20 µM CCCP (plus 3-minute pre-incubation). Saturation of the suramin-receptor interaction was demonstrated by including 100 µM unlabelled suramin (blue bars). Bars represent average and SEM or three independent experiments, each performed in quadruplicate. (C) Correlation plot of pentamidine transport rate versus protonmotive force (PMF), r^2^ = 0.93, P<0.05 by F-test. Concentrations in µM are indicated in the frame. CCCP is shown in red and not included in the regression analysis. Each data point is the average of 4 or more independent repeats performed in quadruplicate. The values for PMF were taken from (De Koning and Jarvis, 1997b). (D) Uptake of 0.25µM [^3^H]-glycerol by *aqp1/aqp2/aqp3* null cells expressing TbAQP2-WT. Dose response with CCCP and pentamidine (PMD), using an incubation time of 1 min. The graph shown was performed in triplicate and representative of three independent repeats. (E) As C but using 0.025 µM [^3^H]-pentamidine and 30 s incubations. Representative graph in triplicate from 3 independent repeats. *, P<0.05; **, P<0.01; ***, P<0.001 by Student’s unpaired t-test.

Although there is a good correlation between the proton-motive force and TbAQP2-mediated pentamidine transport (Fig. 5C), the effect of CCCP was stronger than expected, and stronger than previously observed for [^3^H]-hypoxanthine uptake in *T. brucei* bloodstream forms (De Koning & Jarvis 1997b) and we thus investigated whether CCCP might have a direct effect on TbAQP2. Indeed, CCCP inhibited uptake of (neutral) [^3^H]-glycerol in *tbaqp1-2-3* null cells expressing TbAQP2-WT, with an IC_50_ of 20.7 ± 2.6 µM (n=3) and inhibited [^3^H]-pentamidine uptake in the same cells with a similar IC_50_ (Fig. 5D,E), showing CCCP to inhibit TbAQP2 directly, irrespective of effects on the membrane potential.

### 4. Molecular dynamics modelling of pentamidine interactions with TbAQP2

To further investigate pentamidine binding and permeation in TbAQP2, we used the coordinates of the TbAQP2-pentamidine complex that was modelled in our previous study (Munday *et al,* 2015a). The stability of the protein model was first confirmed by unbiased atomistic molecular dynamics simulations (Supplemental Fig. S6). We then conducted force-probe simulations, in which a moving spring potential was used to enforce unbinding of pentamidine from its docked binding position and subsequently reconstructed the free-energy profile of pentamidine association-dissociation along the pore axis by employing Jarzynski’s equality (Park *et al,* 2003).

Figure 6 shows that the docked position of pentamidine correctly identified its minimum free-energy binding site inside the TbAQP2 pore. Pentamidine adopts an extended state inside the TbAQP2 pore, adapting its molecular shape to the narrow permeation channel; pentamidine binding poses display inter-amidine lengths in the range 16.5 – 17 Å. Importantly, our steered simulations reveal that pentamidine can exit the channel in either direction, and that unbinding on the route towards the cytoplasm occurs on a free-energy surface roughly symmetric to that towards the extracellular side. Apart from overcoming the strongly attractive binding interaction in the centre, there are no major further free-energy barriers in either direction. The computed free-energy profile of pentamidine binding to the TbAQP2 structural model slightly overestimates its experimentally recorded binding affinity. However, the pentamidine conformation binding the narrow pore may not be the lowest-energy internal conformation of the small molecule, a factor that may be underrepresented in the profile as simulations were started from the protein-bound state. A further source of uncertainty stems from the protein model, which is expected to be somewhat less accurate than a crystal structure.

**Fig. 6.**
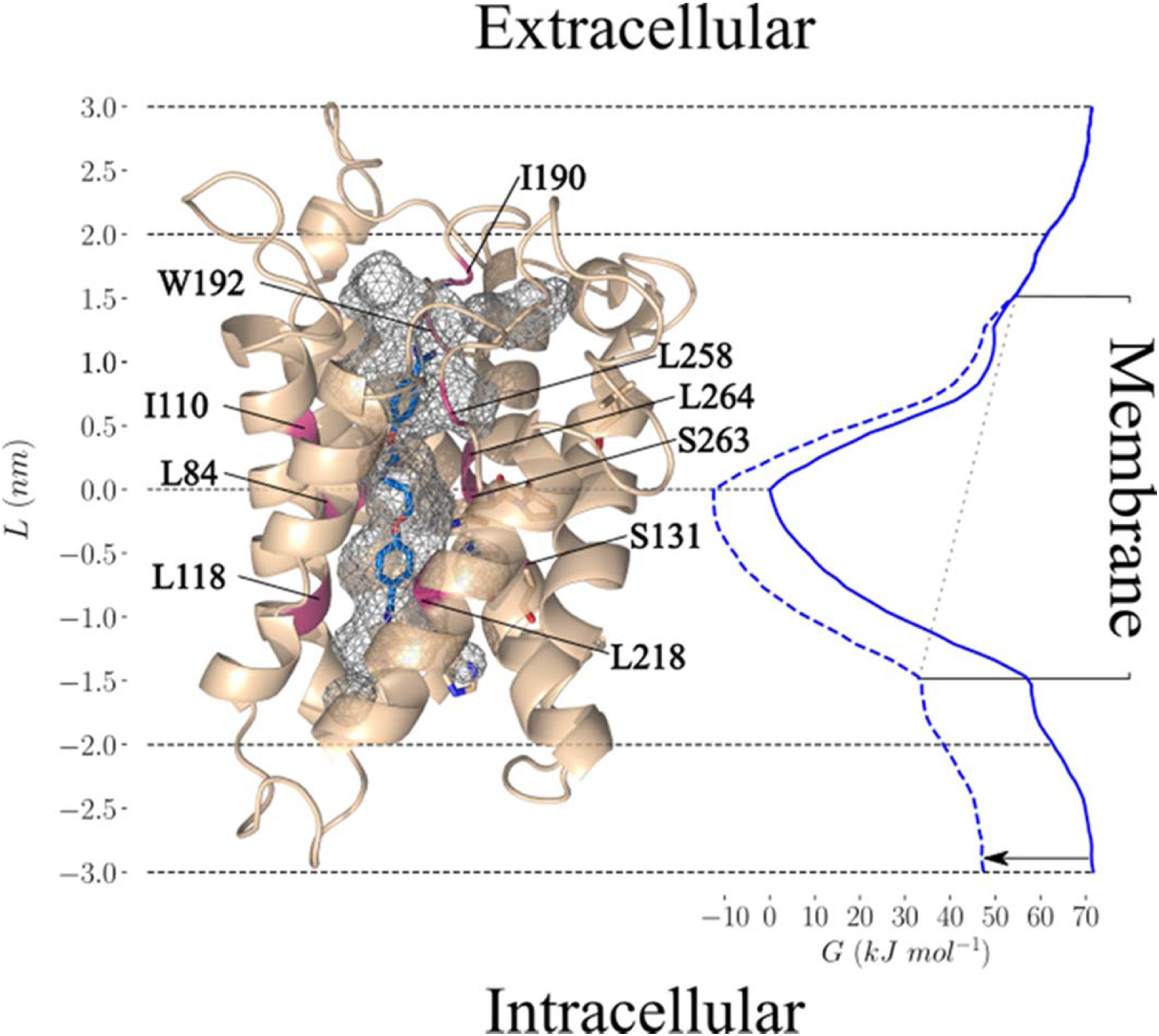
Pentamidine binding in TbAQP2 and free-energy profile of permeation (Left). Docked conformation of pentamidine (blue) bound to the TbAQP2 (wheat). The protein and the ligand were modelled as described.^4^ The protein pore is shown in grey mesh, and the mutated positions described in the text are in magenta. (Right) Free-energy profile G(L) (solid blue line) along the pore axis of TbAQP2 (L). The membrane voltage of *T. b. brucei* gives rise to a voltage drop across the membrane (gray dotted line), which alters the free-energy profile (dashed blue line includes V_m_ effect) and reduces the free-energy of pentamidine exit into the intracellular bulk by ∼22 kJ/mol as compared to the extracellular side (black arrow).

Due to the dicationic character of pentamidine, the free-energy profile of the molecule within TbAQP2 strongly depends on the membrane voltage. The voltage drop of −125 mV across the cytoplasmic membrane of *T. b. brucei* (De Koning & Jarvis, 1997b), with a negative potential inside the cell, results in an overall inward attraction of ∼22 kJ/mol (Fig. 6, arrow), i.e. exit from TbAQP2 into the cytoplasm is substantially more favourable for pentamidine than towards the extracellular side. Taken together, the free-energy profile under membrane voltage explains the strong coupling between pentamidine uptake and V_m_ observed in the experiments. The high affinity of the binding interaction leads to slow off-rates and a relatively low V_max_ (0.0044 ± 0.0004 pmol(10^7^ cells)^-1^s^-1^) (De Koning, 2001a).

### 5. SAR of the pentamidine-AQP2 interaction

In order to study substrate binding and selectivity by the *T. b. brucei* High Affinity Pentamidine Transporter (HAPT1/TbAQP2), competition assays were performed with a series of pentamidine analogues and other potential inhibitors, in the presence of 1 mM unlabelled adenosine to block diamidine uptake by the TbAT1 aminopurine transporter (De Koning, 2001a; Bridges *et al,* 2007). High specific activity [^3^H]-pentamidine was used at 30 nM, below the K_m_ value (De Koning, 2001a). Uptake was linear for at least 3 min (De Koning, 2001a) and we utilized 60 s incubations for the determination of inhibition constants (K_i_). At 30 nM [^3^H]-pentamidine there is virtually no uptake through LAPT1 (Bridges *et al,* 2007) (K_m_ value ∼1000-fold higher than HAPT1) (De Koning, 2001a). The full dataset of 71 compounds is presented in Supplemental Table S1, featuring K_i_s spanning 5 log units.

#### 5.1. The linker length and composition is a strong determinant for high affinity binding of pentamidine

We determined the K_i_ values for analogues with a 2–8 methylene unit linker (Fig. 7A, Table 1). Pentamidine analogues featuring 5-7 units displayed submicromolar binding affinities (5 > 6 > 7), while fewer (3-4) or more (8) only conveyed low micromolar binding affinity, equivalent to a decrease in Gibbs free energy of binding (ΔG^0^) from 10.2 to 13.0 kJ/mol (Table 1). Energy minimalization using Gaussian16 yielded an elongated conformation for pentamidine, with an inter-amidine length of 17.8 Å (Fig. 7B). Replacement of the ether oxygens with S or NH, analogues RT-48 and RT-50, respectively (Table 1) resulted in δ(ΔG^0^) of 10.0 and 12.9 kJ/mol, respectively, indicating that the ether oxygens potentially act as H-bond acceptors: the NH group serves only as an H-bond donor, as its lone pair is conjugated with the aromatic system, and the sulphur mimics an aromatic NH (Beno *et al,* 2015). The sulfone analogue (RT-49), which introduces a dihedral angle of 180° between the benzamidine and the linker (Brameld *et al,* 2008), displayed no binding affinity. We propose that a near-planar conformation of the Phe-O-CH_2_ segment is required for efficient engagement of the binding site. This is supported by examining the binding affinities found for the analogous benzofuramidine series (e.g. RT-14, Fig. 7B), which has a conformationally predefined ether-methylene bond orientation. Replacement of the middle methylene unit of pentamidine with an isosteric oxygen (ethylene glycol derivative DB1699, Table 1) results in a less flexible linker and a remarkable drop in binding affinity (δ(ΔG^0^) = 15.3 kJ/mol).

**Fig. 7.**
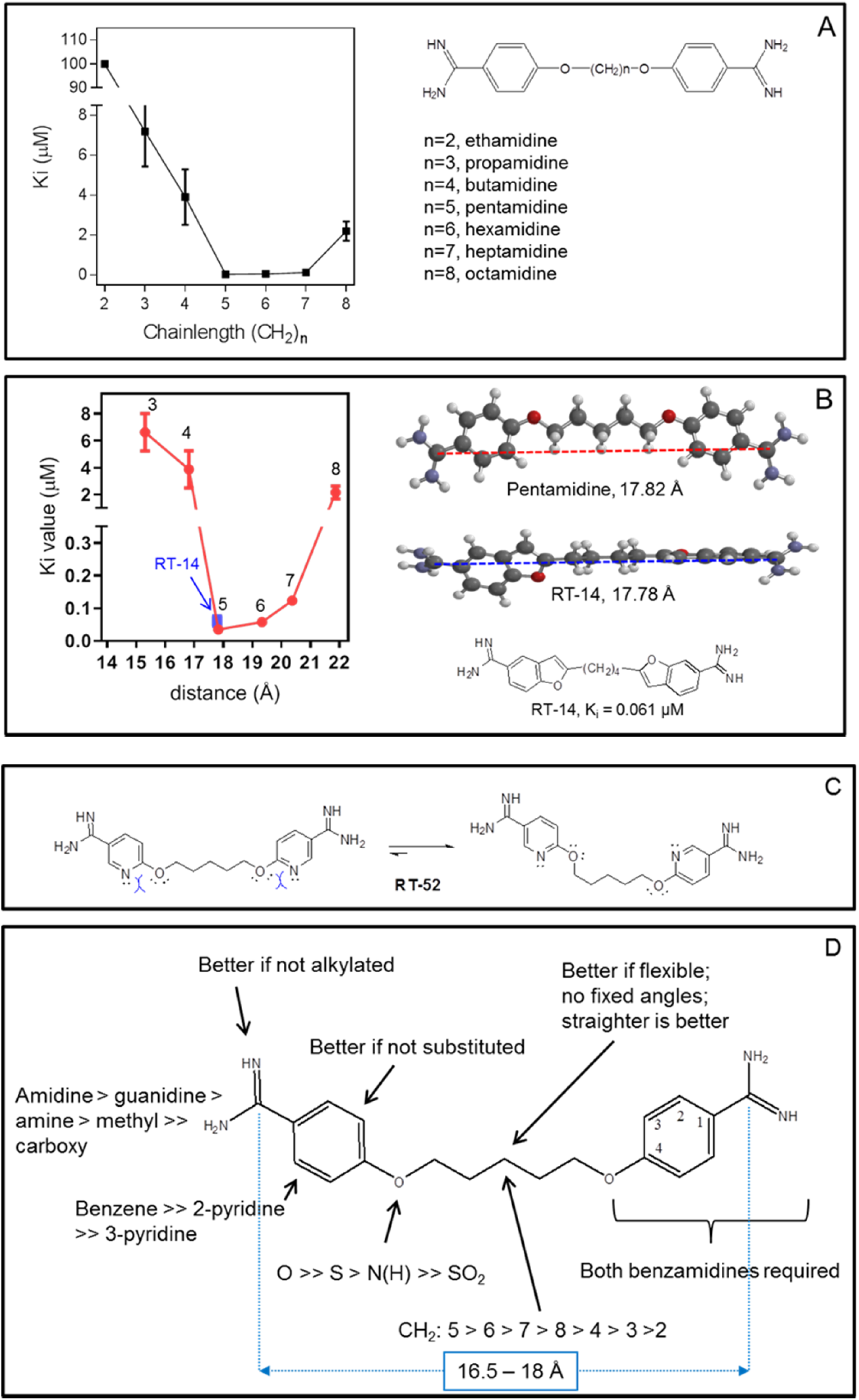
Correlation between linker chain length and affinity to HAPT1. (A) A series of pentamidine analogues with different methylene linker length was tested for inhibition of TbAQP2/HAPT1-mediated 25 nM [^3^H]-pentamidine transport (i.e. in the presence of adenosine to block the TbAT1/P2 transporter). The K_i_ values are listed in Table 1. All K_i_ values are shown as average and SEM of 3 or more independent experiments, each performed in triplicate. (B) The distance between the amidine carbon atoms in the lowest-energy conformation was calculated using density functional theory as implemented in Spartan ’16 v2.0.7. Geometry optimisations were performed with the wB97XD functional and the 6-31G* basis set at the ground state in gas phase. Structures and distances shown represent the dication state that is overwhelmingly prevalent in aqueous solution at neutral pH. The numbered red data points correspond to the propamidine - octamidine series in frame A. (C) Repulsion between free electron pairs (double dots), indicated by curved blue lines for RT-52 in the *cis*-conformation, causing it to exist overwhelmingly in the *anti*-conformation. (D) Overview of SAR observations on the binding preferences of TbAQP2 for pentamidine and its analogues.

**Table 1.**
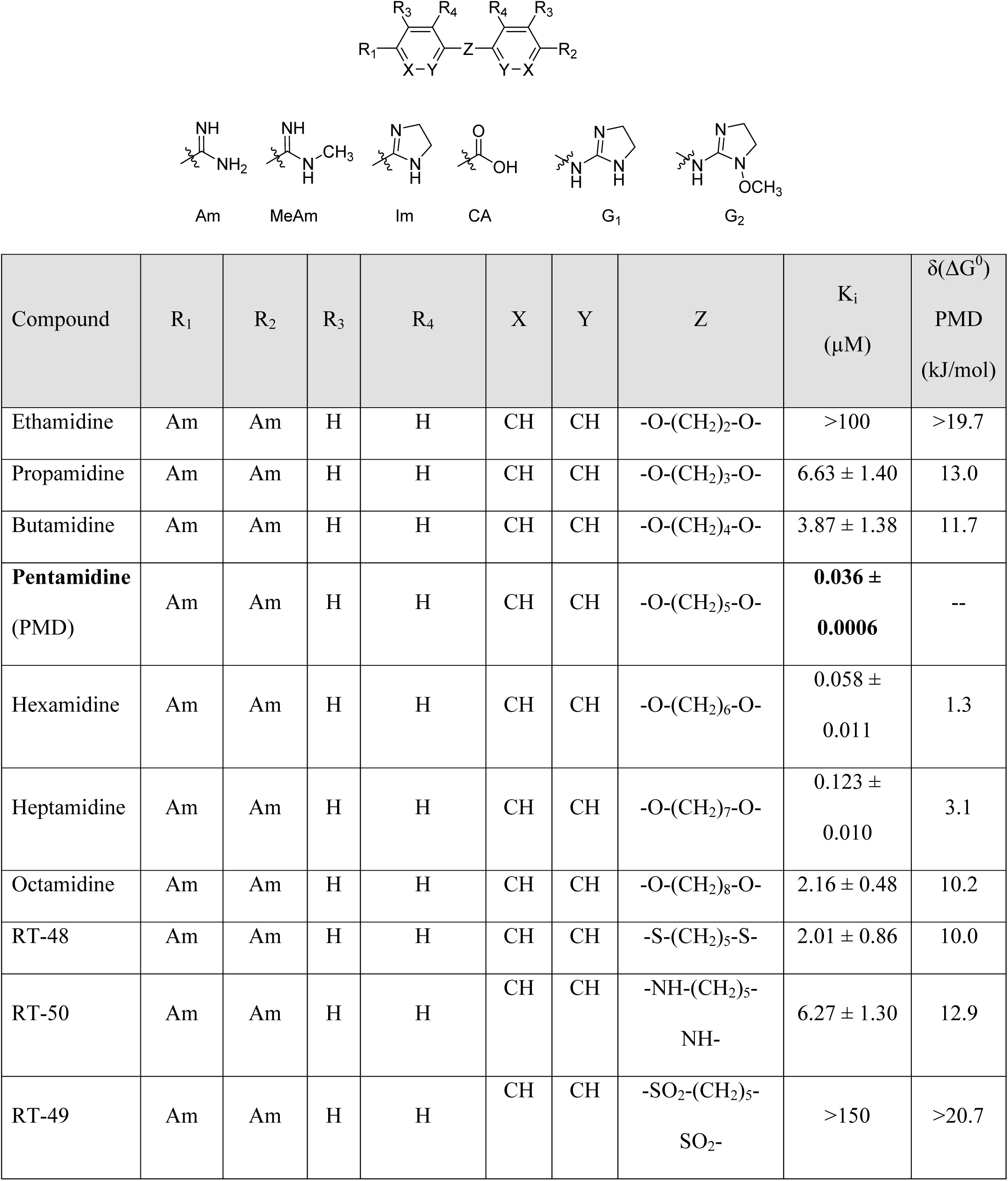
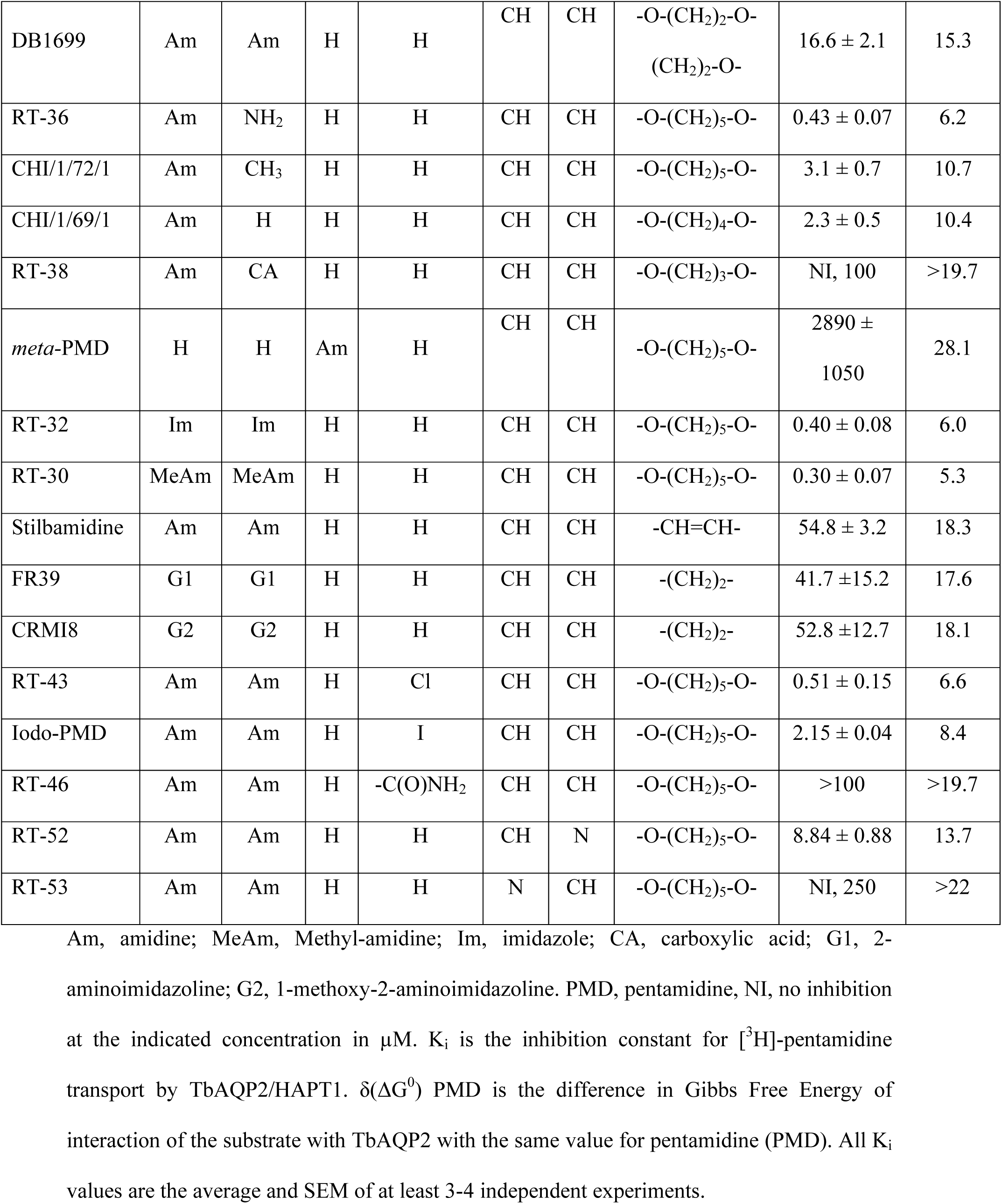
Pentamidine analogues with an aliphatic linker

| Compound | R<sub>1</sub> | R<sub>2</sub> | R<sub>3</sub> | R<sub>4</sub> | X | Y | Z | K<sub>i</sub><br>(μM) | δ(ΔG<sup>0</sup>)<br>PMD<br>(kJ/mol) |
| --- | --- | --- | --- | --- | --- | --- | --- | --- | --- |
| Ethamidine | Am | Am | H | H | CH | CH | -O-(CH<sub>2</sub>)<sub>2</sub>-O- | >100 | >19.7 |
| Propamidine | Am | Am | H | H | CH | CH | -O-(CH<sub>2</sub>)<sub>3</sub>-O- | 6.63 ± 1.40 | 13.0 |
| Butamidine | Am | Am | H | H | CH | CH | -O-(CH<sub>2</sub>)<sub>4</sub>-O- | 3.87 ± 1.38 | 11.7 |
| **Pentamidine**<br>(PMD) | Am | Am | H | H | CH | CH | -O-(CH<sub>2</sub>)<sub>5</sub>-O- | **0.036 ±**<br>**0.0006** | -- |
| Hexamidine | Am | Am | H | H | CH | CH | -O-(CH<sub>2</sub>)<sub>6</sub>-O- | 0.058 ±<br>0.011 | 1.3 |
| Heptamidine | Am | Am | H | H | CH | CH | -O-(CH<sub>2</sub>)<sub>7</sub>-O- | 0.123 ±<br>0.010 | 3.1 |
| Octamidine | Am | Am | H | H | CH | CH | -O-(CH<sub>2</sub>)<sub>8</sub>-O- | 2.16 ± 0.48 | 10.2 |
| RT-48 | Am | Am | H | H | CH | CH | -S-(CH<sub>2</sub>)<sub>5</sub>-S- | 2.01 ± 0.86 | 10.0 |
| RT-50 | Am | Am | H | H | CH | CH | -NH-(CH<sub>2</sub>)<sub>5</sub>-<br>NH- | 6.27 ± 1.30 | 12.9 |
| RT-49 | Am | Am | H | H | CH | CH | -SO<sub>2</sub>-(CH<sub>2</sub>)<sub>5</sub>-<br>SO<sub>2</sub>- | >150 | >20.7 |
| DB1699 | Am | Am | H | H | CH | CH | -O-(CH<sub>2</sub>)<sub>2</sub>-O-(CH<sub>2</sub>)<sub>2</sub>-O- | 16.6 ± 2.1 | 15.3 |
| RT-36 | Am | NH<sub>2</sub> | H | H | CH | CH | -O-(CH<sub>2</sub>)<sub>5</sub>-O- | 0.43 ± 0.07 | 6.2 |
| CHI/1/72/1 | Am | CH<sub>3</sub> | H | H | CH | CH | -O-(CH<sub>2</sub>)<sub>5</sub>-O- | 3.1 ± 0.7 | 10.7 |
| CHI/1/69/1 | Am | H | H | H | CH | CH | -O-(CH<sub>2</sub>)<sub>4</sub>-O- | 2.3 ± 0.5 | 10.4 |
| RT-38 | Am | CA | H | H | CH | CH | -O-(CH<sub>2</sub>)<sub>3</sub>-O- | NI, 100 | >19.7 |
| *meta*-PMD | H | H | Am | H | CH | CH | -O-(CH<sub>2</sub>)<sub>5</sub>-O- | 2890 ± 1050 | 28.1 |
| RT-32 | Im | Im | H | H | CH | CH | -O-(CH<sub>2</sub>)<sub>5</sub>-O- | 0.40 ± 0.08 | 6.0 |
| RT-30 | MeAm | MeAm | H | H | CH | CH | -O-(CH<sub>2</sub>)<sub>5</sub>-O- | 0.30 ± 0.07 | 5.3 |
| Stilbamidine | Am | Am | H | H | CH | CH | -CH=CH- | 54.8 ± 3.2 | 18.3 |
| FR39 | G1 | G1 | H | H | CH | CH | -(CH<sub>2</sub>)<sub>2</sub>- | 41.7 ± 15.2 | 17.6 |
| CRMI8 | G2 | G2 | H | H | CH | CH | -(CH<sub>2</sub>)<sub>2</sub>- | 52.8 ± 12.7 | 18.1 |
| RT-43 | Am | Am | H | Cl | CH | CH | -O-(CH<sub>2</sub>)<sub>5</sub>-O- | 0.51 ± 0.15 | 6.6 |
| Iodo-PMD | Am | Am | H | I | CH | CH | -O-(CH<sub>2</sub>)<sub>5</sub>-O- | 2.15 ± 0.04 | 8.4 |
| RT-46 | Am | Am | H | -C(O)NH<sub>2</sub> | CH | CH | -O-(CH<sub>2</sub>)<sub>5</sub>-O- | >100 | >19.7 |
| RT-52 | Am | Am | H | H | CH | N | -O-(CH<sub>2</sub>)<sub>5</sub>-O- | 8.84 ± 0.88 | 13.7 |
| RT-53 | Am | Am | H | H | N | CH | -O-(CH<sub>2</sub>)<sub>5</sub>-O- | NI, 250 | >22 |

Am, amidine; MeAm, Methyl-amidine; Im, imidazole; CA, carboxylic acid; G1, 2- aminoimidazoline; G2, 1-methoxy-2-aminoimidazoline. PMD, pentamidine, NI, no inhibition at the indicated concentration in $\mu\text{M}$ . $K_i$ is the inhibition constant for [<sup>3</sup>H]-pentamidine transport by TbAQP2/HAPT1. $\delta(\Delta G^0)$ PMD is the difference in Gibbs Free Energy of interaction of the substrate with TbAQP2 with the same value for pentamidine (PMD). All $K_i$ values are the average and SEM of at least 3-4 independent experiments.

#### 5.2. Two amidine groups are required for high affinity binding

Matched-molecular pair analysis of non-symmetric analogues identified that both amidines contribute to high affinity binding (compare pairs pentamidine/RT-36 and pentamidine/CHI/1/72/1; Table 1). Removal of an H-bond donor (as in CHI/1/72/1) leads to a loss in ΔG^0^ >10 kJ/mol. The aniline derivative RT-36 can still act as an H-bond donor, albeit with significantly reduced basicity (and thus H-bond acceptor propensity), and accordingly displayed intermediate affinity (δ(ΔG^0^) = 6.2 kJ/mol). Interestingly, the removal of one amidine (compare butamidine and CHI/1/69/1) did not produce a significant effect on the binding affinity (K_i_ = 3.87 µM and K_i_ = 2.33 µM, respectively), indicating that the low affinity of butamidine (compare 36 nM for pentamidine) is due to an inability to attain a productive interaction with the second amidine. Capping of the amidine group, resulting in imidazoline analogue RT-32, or methylation (analogue RT-30) reduced binding to HAPT1, probably due to increased steric crowding at the interaction site, impairing H-bonding. Reducing pentamidine to just 4-hydroxybenzamidine removed essentially all affinity (K_i_ = 2.9 mM; δ(ΔG^0^) = 28.1 kJ/mol), and the replacement of one amidine with a carboxylic group (compare propamidine, RT-38) was highly deleterious for engagement with the binding site. Finally, the orientation of the amidine group is crucial as shown by a *meta* to *para* change on the phenyl ring (*meta*-pentamidine, Table 1). We conclude that for high affinity both amidine groups must be able to interact unimpeded with the transporter, and in the linear (*para*) conformation.

#### 5.3. Fully conjugated linking units

Stilbamidine and the short-linker analogues FR39 and CRMI8 (Ríos Martínez *et al,* 2015) displayed low binding affinity (Table 1). Diminazene also displayed similar low affinity (K_i_ = 63 µM), and [^3^H]-diminazene uptake can only just be detected in procyclic *T. b. brucei*, *i.e.* in the absence of the TbAT1/P2 transporter (Teka *et al,* 2011), potentially indicating a minimal uptake via HAPT1. Stilbamidine and diminazene feature a similar inter-amidine distance, much shorter than pentamidine (12.35 and 12.25 Å, respectively). DB75 (furamidine) likewise displayed low affinity (Table 2) and is only internalised by TbAT1/P2 (Ward *et al,* 2011). The 2,5-furan linker imposes a fixed, inflexible angle of 136° on the benzamidine moieties and the phenyl rings will adopt a planar orientation with respect to the furan plane. This appears to allow only one benzamidine end to interact with the transporter, as (unlike the flexible linker of aliphatic diamidines, *vide supra*) the replacement of one amidine group actually increases the binding affinity, presumably by allowing an improved bonding orientation of the remaining amidine. Thus, DB607 (methoxy for amidine) and DB960 (*N*-methyl benzimidazole for benzamidine) display a somewhat higher affinity than DB75, although the fixed angle was unchanged. Introduction of a pyridine-N in the *ortho*-position with respect to the amidine functionality (DB994), dramatically reduces the pK_a_ of the amidine moiety (Wang *et al,* 2010), resulting in a complete loss of binding affinity (K_i_ = 167 ± 20 µM), while this was not observed for the corresponding *meta*-pyridine derivative (DB829). The unfavourable furan bond angle is further demonstrated by the distally elongated analogues DB1061 and DB1062 that approximate the inter-amidine distance of pentamidine but showed no improvement in binding affinity (Table 2). Replacement of furan with thiazole (ER1004) or methylpyrrole (DB320), which feature a similar bond angle, also revealed comparable binding affinities. In contrast, a 2,5-substituted thiophene (DB686 and DB1063) or 2,5-substituted selenophene (DB1213) as a bio-isosteric replacement for the furan linker resulted in significantly higher binding affinities when compared to their matched pair analogue (DB1063/DB1061 and DB1213/DB75), which we attribute to a larger benzamidine-benzamidine angle. This is corroborated by the much weaker binding of the 2,4-thiophene derivative DB1077. A terminal amidine cap (imidazoline) reduced affinity as it did for pentamidine (compare DB1061/DB1062 and DB1063/DB1064 (Table 2)). A difuran spacer (DB914) resulted in a high affinity binder (K_i_ = 0.073 µM) because the two furans orient themselves in a *trans* conformation, resulting in a near-linear molecule.

**Table 2.**
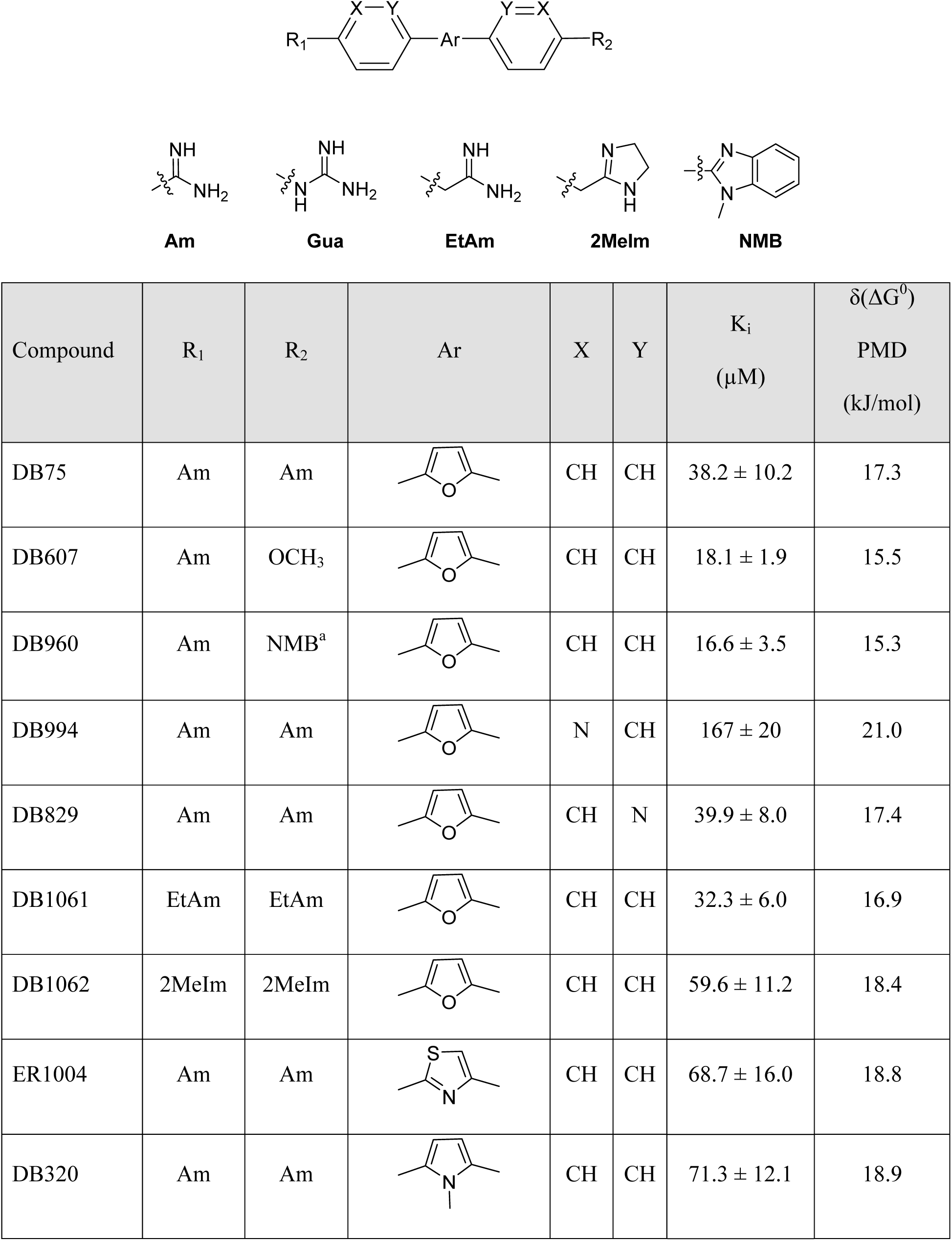
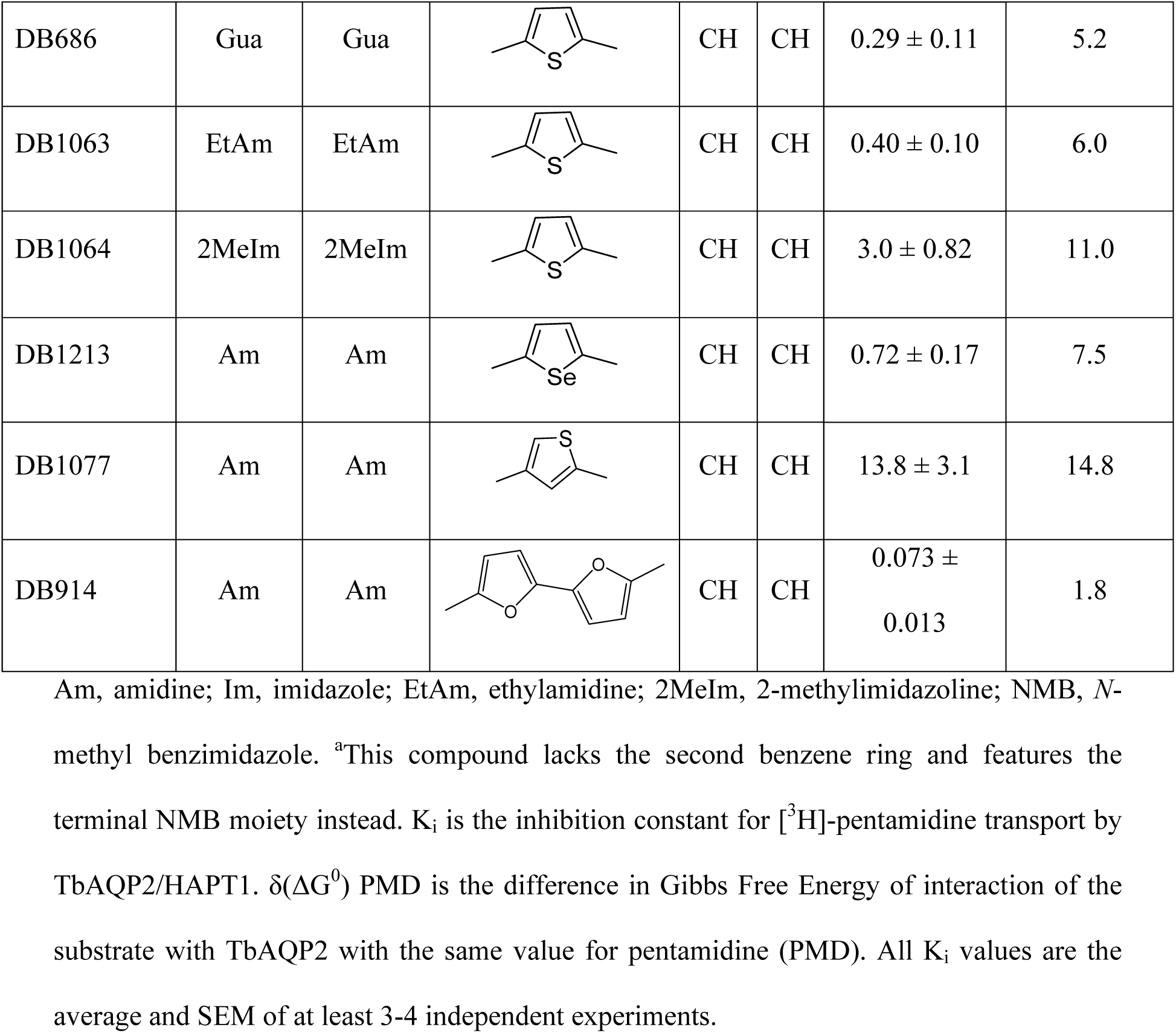
Selection of diamidine analogues with aromatic linkers

| Compound | R<sub>1</sub> | R<sub>2</sub> | Ar | X | Y | K<sub>i</sub><br>(μM) | δ(ΔG<sup>0</sup>)<br>PMD<br>(kJ/mol) |
| --- | --- | --- | --- | --- | --- | --- | --- |
| DB75 | Am | Am |  | CH | CH | 38.2 ± 10.2 | 17.3 |
| DB607 | Am | OCH<sub>3</sub> |  | CH | CH | 18.1 ± 1.9 | 15.5 |
| DB960 | Am | NMB<sup>a</sup> |  | CH | CH | 16.6 ± 3.5 | 15.3 |
| DB994 | Am | Am |  | N | CH | 167 ± 20 | 21.0 |
| DB829 | Am | Am |  | CH | N | 39.9 ± 8.0 | 17.4 |
| DB1061 | EtAm | EtAm |  | CH | CH | 32.3 ± 6.0 | 16.9 |
| DB1062 | 2MeIm | 2MeIm |  | CH | CH | 59.6 ± 11.2 | 18.4 |
| ER1004 | Am | Am |  | CH | CH | 68.7 ± 16.0 | 18.8 |
| DB320 | Am | Am |  | CH | CH | 71.3 ± 12.1 | 18.9 |
| DB686 | Gua | Gua | | CH | CH | $0.29 \pm 0.11$ | 5.2 |
| DB1063 | EtAm | EtAm | | CH | CH | $0.40 \pm 0.10$ | 6.0 |
| DB1064 | 2MeIm | 2MeIm | | CH | CH | $3.0 \pm 0.82$ | 11.0 |
| DB1213 | Am | Am | | CH | CH | $0.72 \pm 0.17$ | 7.5 |
| DB1077 | Am | Am | | CH | CH | $13.8 \pm 3.1$ | 14.8 |
| DB914 | Am | Am | | CH | CH | $0.073 \pm 0.013$ | 1.8 |

Am, amidine; Im, imidazole; EtAm, ethylamidine; 2MeIm, 2-methylimidazoline; NMB, *N*- methyl benzimidazole. <sup>a</sup>This compound lacks the second benzene ring and features the terminal NMB moiety instead. $K_i$ is the inhibition constant for [<sup>3</sup>H]-pentamidine transport by TbAQP2/HAPT1. $\delta(\Delta G^0)$ PMD is the difference in Gibbs Free Energy of interaction of the substrate with TbAQP2 with the same value for pentamidine (PMD). All $K_i$ values are the average and SEM of at least 3-4 independent experiments.

#### 5.4. Modifications to the phenyl rings of pentamidine

Substituents in the *ortho*-position (relative to the alkyloxy substituent) of pentamidine were poorly tolerated, including chloride or iodide (RT-43, iodopentamidine; Table 1); the amide analogue displayed no affinity at all (RT-46). Such substituents will cause an out-of-plane conformation of the alkoxy-group to avoid clashing with the *ortho*-substituent; high-affinity pentamidine binding appears to require a coplanar arrangement of the first methylene bound to the oxygen. Similarly, the introduction of an *ortho*-pyridine N (RT-52) led to a δ(ΔG^0^) of 13.7 kJ/mol. This derivative exhibits a conformational bias towards an *anti*-orientation of the ether oxygen and pyridine nitrogen (Fig. 7C) (Chein & Corey, 2010). The regio-isomeric *meta*-pyridine (RT-53) was completely inactive, reflecting the need for a positively charged amidine, as this analogue has a significantly reduced pKa (Wang *et al,* 2010) (see furan-spaced analogue DB994, *supra*).

#### 5.5. Non-diamidine trypanocides

The important veterinary trypanocide isometamidium, a hybrid of the phenanthridine ethidium and the diamidine diminazene, inhibited HAPT1-mediated [^3^H]-pentamidine uptake with a K_i_ of only 3.5 µM (Supplemental Table S1), most probably through an interaction with its benzamidine moiety, as ethidium displayed virtually no affinity (K_i_ = 97 µM). However, we found no evidence that HAPT1/AQP2 is able to transport the bulky isometamidium molecule. For instance, the 2T1, *tbaqp2* null, TbAQP2 expressed in *tbaqp2* null, and the *tbaqp2/tbaqp3* null strains displayed statistically identical EC_50_ values for isometamidium (112 ± 12 nM, 103 ± 14 nM, 98 ± 24 nM and 95 ± 12 nM, respectively; P>0.05, Student’s unpaired t-test), and the EC_50_ values for ethidium were also identical for each of these strains (1.32 ± 0.07 µM, 1.39 ± 0.08 µM, 1.35 ± 0.11 µM and 1.38 ± 0.14 µM, respectively). It is thus likely that isometamidium acts as an extracellular inhibitor rather than a substrate for HAPT/AQP2, as it does for the TbAT1/P2 transporter (De Koning, 2001b). The nitro-heterocyclic trypanocide megazol (Carvalho *et al,* 2014), curcumin and its trypanocidal analogue AS-HK14 (Alkhaldi *et al,* 2015) failed to inhibit HAPT1. Two trypanocidal bis-phosphonium compounds, CD38 (Taladriz *et al,* 2012) and AHI43 (Alkhaldi *et al,* 2016) did inhibit pentamidine uptake (K_i_ 5-10 µM), whereas two related compounds, CDIV31 and AHI15 (Taladriz *et al,* 2012), did not. Phloretin, which inhibits human AQP9 and AQP3 (Geng *et al,* 2017),^47^ displayed a K_i_ of 1.76 µM for HAPT1/TbAQP2.

#### 5.6. Are all the HAPT1/AQP2 inhibitors transported?

In an uptake-by-endocytosis model it would be expected that TbAQP2 binding energy correlates well with TbAQP2-mediated uptake rates for each analogue. We were unable to ascertain the existence of such a correlation directly, for lack of radiolabelled substrates other than pentamidine and diminazene and thus used the Resistance Factor (RF; EC_50_(aqp2/3 null)/EC_50_(TbAQP2-WT)) as a proxy: clearly, a compound with a significant RF is internalized by TbAQP2. We observed a poor correlation between HAPT1 binding affinity and the level of resistance in the *tbaqp2/tbaqp3* null strain (r^2^=0.039, Supplemental Fig. S7; n=30), with many inhibitors, even those with high affinity, not displaying any significant resistance in the null line. This indicates that many of these inhibitors inhibit HAPT1/TbAQP2 but are not transported by it. This is not compatible with a model in which pentamidine binds and is then internalized by endocytosis: the inhibitors do not show resistance in the *tbaqp2/tbaqp3* null line, whereas substrates do. The caveat inherent to using the RF instead of rate of transport is that it cannot be excluded that some of the test compounds are AQP2 substrates yet predominantly taken up by transporters other than TbAQP2, and hence show a low RF.

#### 5.7. SAR summary

Figure 7D summarises the structure-activity relationship of pentamidine interactions with HAPT1/TbAQP2. No modification in any part of the molecule improved affinity for TbAQP2, but virtually every modification resulted in a significant loss of binding activity. The results clearly demonstrate that at least both amidine groups and one or both ether oxygens are involved in interactions with AQP2, the sum of which adds up to the unusually high binding energy for this substrate-transporter pair (ΔG^0^ = −42.6 kJ/mol). These results, unambiguously demonstrating pentamidine binding in an elongated orientation, are in complete agreement with the modelling and molecular dynamics, and the mutational analysis presented above, strengthening those conclusions using a completely different approach.

## Discussion and conclusion

There is overwhelming consensus that expression of TbAQP2 is associated with the extraordinary sensitivity of *T. brucei* to pentamidine and melaminophenyl arsenicals, and that mutations and deletions in this locus cause resistance (Baker *et al,* 2012, 2013; Graf *et al,* 2013, 2015, 2016; Pyana Pati *et al,* 2014; Munday *et al,* 2014, 2015a; Unciti-Broceta *et al,* 2015). What has remained however unclear is the mechanism underpinning these phenomena – there are currently no documented other examples of aquaporins transporting such large molecules.. Yet, considering how ubiquitous aquaporins are to almost all cell types, this question is of wide pharmacological importance: if large cationic and neutral drugs (pentamidine and melarsoprol, respectively) can be taken up via an aquaglyceroporin of *T. brucei*, what other pharmacological or toxicological roles may these channels be capable of in other cell types? This manuscript shows clearly that changes in the TbAQP2 WGYR and NPA/NPA motifs, which collectively enlarge the pore and remove the cation filter, allow the passage of these drugs into the cell, and thereby underpin the very high sensitivity of the parasite to these drugs.

TbAQP2 has evolved, apparently by positive selection given the high dN/dS ratio, to remove all main constriction points, including the aromatic amino acids and the cationic arginine of the selectivity filter, and the NPA/NPA motif, resulting in an unprecedentedly enlarged pore size. Whereas the advantage of this to *T. b. brucei* is yet unknown, the adaptation is stable within the *brucei* group of trypanosomes, and found in *T. b. rhodesiense* (Munday *et al,* 2014; Graf *et al,* 2016), *T. b. gambiense* (Graf *et al,* 2013, 2015; Munday *et al,* 2014; Pyana Pati *et al,* 2014), *T. equiperdum* and *T. evansi* (Philippe Büscher and Nick Van Reet, unpublished). As such, it is not inappropriate to speculate that the wider pore of TbAQP2 (i) allows the passage of something not transported by TbAQP1 and TbAQP3; (ii) that this confers an a yet unknown advantage to the cell; and (iii) that uptake of pentamidine is a by-product of this adaptation.

It is difficult to reconcile the literature on pentamidine transport/resistance with uptake via endocytosis. For instance, the rate of endocytosis in bloodstream trypanosomes is much higher than in the procyclic lifecycle forms (Langreth & Balber, 1975; Zoltner *et al,* 2016), yet the rate of HAPT-mediated [^3^H]-pentamidine uptake in procyclics is ∼10-fold higher than in bloodstream forms (De Koning, 2001a; Teka *et al,* 2011), despite the level of TbAQP2 expression being similar in both cases (Siegel *et al,* 2010; Jensen *et al,* 2014). Moreover, in procyclic cells TbAQP2 is spread out over the cell surface (Baker *et al,* 2012) but endocytosis happens exclusively in the flagellar pocket (Field & Carrington, 2009) (which is 3-fold smaller in procyclic than in bloodstream forms (Demmel *et al,* 2014)), as the pellicular microtubule networks below the plasma membrane prevent endocytosis (Zoltner *et al,* 2016). Similarly, the expression of TbAQP2 in *Leishmania mexicana* promastigotes produced a rate of [^3^H]-pentamidine uptake more than 10-fold higher than observed in *T. brucei* BSF (Munday *et al,* 2014), despite these cells also having a low endocytosis rate (Langreth & Balber, 1975). The K_m_ and inhibitor profile of the TbAQP2-mediated pentamidine transport in these promastigotes was indistinguishable from HAPT in procyclic or bloodstream form *T. brucei* (De Koning, 2001a). Moreover, the experimental V_max_ for HAPT-mediated pentamidine uptake in *T. brucei* BSF and procyclics (Baker *et al,* 2013) can be expressed as 9.5×10^5^ and 8.5×10^6^ molecules/cell/h, respectively; given a 1:1 stoichiometry for AQP2:pentamidine the endocytosis model would require the internalisation and recycling of as many units of TbAQP2 and this seems unlikely, especially in procyclic cells. These observations are all inconsistent with the contention that pentamidine uptake by trypanosomes is dependent on endocytosis.

Furthermore, the Gibbs free energy of −42 kJ/mol for the pentamidine/AQP2 interaction (De Koning, 2001a; Zoltner *et al,* 2016) is highly unlikely to be the result of the one interaction between one terminal amidine and Asp265 as required in the endocytosis model (Song *et al,* 2016). For the TbAT1 transporter, a double H-bond interaction of Asp140 with the N1(H)/C(6)NH_2_ motif of adenosine or with one amidine of pentamidine (Munday *et al,* 2015b) is estimated to contribute only ∼16 kJ/mol to the total ΔG^0^ of −34.5 kJ/mol for adenosine (−36.7 kJ/mol, pentamidine) (De Koning & Jarvis, 1999). The endocytosis model also does not address the internalisation of melaminophenyl arsenicals, which presumably would equally need access to Asp265.

Here we systematically mapped the interactions between the aquaporin and pentamidine (ΔG^0^ for 71 compounds), yielding a completely consistent SAR with multiple substrate-transporter interactions, summarised in Fig. 7D. The evidence overwhelmingly supports the notion that pentamidine engages TbAQP2 with both benzamidine groups and most probably with at least one of the linker oxygens, and that its flexibility and small width are both required to optimally interact with the protein. This is completely corroborated by molecular dynamics modelling, which shows minimal energy to be associated with a near-elongated pentamidine centrally in the TbAQP2 pore, without major energy barriers to exiting in either direction, but driven to the cytoplasmic side by the membrane potential. This contrasts with the contention (Song *et al,* 2016) that pentamidine could not be a permeant for TbAQP2 because it did not transport some small cations and that this proves that the larger pentamidine cannot be a substrate either. There is scant rationale for that assertion: out of many possible examples: there are 5 orders of magnitude difference in affinity for pentamidine and *para*-hydroxybenzamidine (35 nM *vs* 2.9 mM; Supplemental Table S1); adenine is not a substrate for the *T. brucei* P1 adenosine transporter (De Koning & Jarvis, 1999), the SLC1A4 and SLC1A5 neutral amino acid transporters transport Ala, Ser, Cys and Thr but not Gly (Kania *et al,* 2013), and Na^+^ is not a permeant of K^+^ channels (Zhorov & Tikhonov, 2013).

The endocytosis model identifies only two key residues for pentamidine access (Leu264) and binding (Asp265) in TbAQP2 (Song *et al,* 2016). Yet, multiple clinical isolates and laboratory strains contain chimeric AQP2/3 genes associated with resistance and/or non-cure that have retained those residues and should thus allow binding and internalisation of pentamidine (Graf *et al,* 2013; Pyana Pati, 2014; Unciti Broceta *et al,* 2015; Munday *et al,* 2014). Although we find that introduction of the AQP3 Arg residue in position 264 (TbAQP2^L264R^) disables pentamidine transport, this is because the positively charged arginine, in the middle of the pore, is blocking the traversing of all cations through the pore, as is its common function in aquaporins (Beitz *et al,* 2006; Wu *et al,* 2009). Indeed, the W(G)YR filter residues appear to be key determinants for pentamidine transport by AQPs and the introduction of all three TbAQP2 residues into TbAQP3 (AQP3^W102I/R256L/Y250L^) was required to create an AQP3 that at least mildly sensitised to pentamidine, and facilitated a detectable level of pentamidine uptake. Conversely, any one of the mutations I110W, L258Y or L264R was sufficient to all but abolish pentamidine transport by TbAQP2. Similarly, the conserved NPA/NPA motif, and particularly the Asp residues, present in TbAQP3 but N<u>S</u>A/NP<u>S</u> in TbAQP2, is also associated with blocking the passage of cations (Wree *et al,* 2011). The unique serine residues in this TbAQP2 motif, halfway down the pore, might be able to make hydrogen bonds with pentamidine. Reinstating the NPA/NPA motif resulted in a TbAQP2 variant with a 93.5% reduced rate of [^3^H]-pentamidine transport.

Tryptophan residues were introduced towards the cytoplasmic end of the TbAQP2 pore (L84W, L118W, L218W) to test the hypothesis that introducing bulky amino acids in that position would block the passage of pentamidine. Each of these mutants was associated with reduced sensitivity to pentamidine and cymelarsan and a >90% reduction in [^3^H]-pentamidine uptake. This effect was size-dependent as the pentamidine transport rate of L84M and L218M was statistically identical to that of control TbAQP2 cells, and L118M also displayed a higher transport rate than L118W (P<0.0001). These mutant AQPs were still functional aquaglyceroporins as their expression in *tbaqp1-2-3* null cells made those cells less sensitive to the TAO inhibitor SHAM, and increased glycerol uptake.

Independence from endocytosis was demonstrated by employing the tetracycline-inducible CRK12 RNAi cell line previously described to give a highly reproducible and progressive endocytosis defect in *T. brucei* (Monnerat *et al,* 2013), and unambiguously distinguishes between uptake via endocytosis and transporters. Twelve hours after CRK12 RNAi induction pentamidine transport was not significantly reduced although uptake of [^3^H]-suramin, which is accumulated by endocytosis through the *T. brucei* flagellar pocket (Zoltner *et al,* 2016), was reduced by 33% (P=0.0027), indicating successful timing of the experiment to the early stage of endocytosis slow-down. Although several ionophores, including CCCP, nigericin and gramicidin strongly inhibited pentamidine uptake, similar to what has been previously reported for transport processes in *T. brucei* that are linked to the protonmotive force (De Koning & Jarvis, 1997a,b, 1998; De Koning *et al,* 1998), this is probably due to the inside-negative membrane potential of −125 mV (De Koning & Jarvis, 1997b) attracting the dicationic pentamidine. This is consistent with the prediction of the molecular dynamics modelling, and the reported role of the HA1–3 proton pumps in pentamidine but not melarsoprol resistance (Alsford *et al,* 2012; Baker *et al,* 2013).

Altogether, we conclude that the primary entry of the sleeping sickness drugs pentamidine and melarsoprol into *T. brucei* spp. is through the unusually large pore of TbAQP2, rendering the parasite extraordinarily sensitive to the drugs (compare *Leishmania mexicana* (Munday *et al,* 2014)). This is the first report providing detailed mechanistic evidence of the uptake of organic drugs (of MW 340 and 398, respectively) by an aquaporin. We show that this porin has evolved through positive selection and identify the adaptations in the constriction motifs that enabled it. We consider that other pore-opening adaptations may have evolved in other organisms, including pathogens, which could initiate the pharmacological exploitation of aquaporins and lead to the design of new drug delivery strategies.

## Materials and Methods

### Trypanosome strains and cultures

The drug-sensitive clonal *T. b. brucei* strain 427 (MiTat 1.2/BS221) (De Koning *et al,* 2000) was used for all the work on the SAR of pentamidine transport. The *tbaqp2/tbaqp3* null cells (Baker *et al,* 2012) and *tbaqp1-2-3* null cells (Jeacock *et al,* 2017) (both obtained from David Horn, University of Dundee, UK) are derived from the 2T1 strain of *T. b. brucei* (Alsford & Horn, 2008). The *CRK12* RNAi cell line^28^ was obtained from Dr Tansy Hammarton (University of Glasgow, UK) and is also based on the 2T1 cell line; RNAi expression was induced with 1 µg/ml tetracycline in the medium. All experiments were performed with bloodstream form trypanosomes grown *in vitro* in HMI-11 medium as described (Wallace *et al,* 2002) at 37 °C in a 5% CO_2_ atmosphere. Cultures were routinely maintained in 10 ml of this medium, being seeded at 5×10^4^ cells/ml and passed to fresh medium at reaching approximately 3×10^6^ cells/ml after 48 h. For transport experiments 150 or 200 ml of culture was seeded at the same density in large flasks and incubated until the culture reached late-log phase.

### Materials

A complete list of diamidine analogues and other chemicals used for the SAR study is given as a table in the supplemental materials with their sources (Supplemental Table S1). Ionophores and uncouplers nigericin, gramicidin, carbonyl cyanide m-chlorophenyl hydrazone (CCCP) and valinomycin, as well as the *T. brucei* proton pump inhibitor *N*-ethylmaleimide (NEM) were all purchased from Sigma-Aldrich. New compounds synthesised for this study are listed and described in the Supplemental Materials.

*Transport of [^3^H]-pentamidine* - Transport of [^3^H]-pentamidine was performed exactly as previously described for various permeants (Wallace *et al,* 2002; Bridges *et al,* 2007; Teka *et al,* 2011) in a defined assay buffer (AB; 33 mM HEPES, 98 mM NaCl, 4.6 mM KCl, 0.55 mM CaCl_2_, 0.07 mM MgSO_4_, 5.8 mM NaH_2_PO_4_, 0.3 mM MgCl_2_, 23 mM NaHCO_3_, 14 mM glucose, pH 7.3). [^3^H]-pentamidine was custom-made by GE Healthcare Life Sciences (Cardiff, UK) with a specific activity of 88 Ci/mmol. Incubations of bloodstream form trypanosomes with 30 nM of this label (unless otherwise indicated) were performed in AB at room temperature for 60 s (unless otherwise indicated) and terminated by addition of 1 ml ice-cold ‘stop’ solution (1 mM unlabelled pentamidine (Sigma) in AB) and immediate centrifugation through oil (7:1 dibutylphthalate:mineral oil v/v (both from Sigma)). Transport was assessed in the presence of 1 mM adenosine to block uptake through the P2 aminopurine transporter; adenosine does not affect HAPT1-mediated transport (De Koning, 2001a; Bridges *et al,* 2007). Inhibition assays were performed routinely with 6 - 10 different concentrations of inhibitor over the relevant range, diluting stepwise by one third each time, in order to obtain a well-defined and accurate sigmoid plot and IC_50_ value (inhibitor concentration giving 50% inhibition of pentamidine transport; calculated by non-linear regression using Prism 6.0 (GraphPad), using the equation for a sigmoid curve with variable slope). Highest concentration was usually 1 mM unless this was shown to be insufficient for good inhibition, or when limited by solubility. *K*_i_ values were obtained from IC_50_ values using

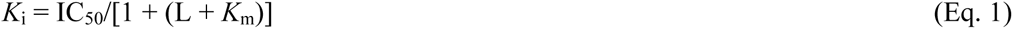

in which L is the [^3^H]-pentamidine concentration and K_m_ the Michaelis-Menten constant for pentamidine uptake by HAPT1 (Wallace *et al,* 2002). The Gibbs Free energy of interaction ΔG^0^ was calculated from

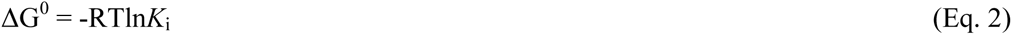

in which R is the gas constant and T is the absolute temperature (Wallace *et al,* 2002).

### Construction of AQP mutants and transfection

All mutations in the TbAQP2 and TbAQP3 genes were introduced to the relevant backbone WT vector, either pRPa^GFP-AQP2^ or pRPa^GFP-AQP3^ (Baker *et al,* 2012), by site-directed mutagenesis. For mutations S131P, S263A, I110W, L264R, L258Y, I190T and W192G in AQP2, and W102I, R256L and Y250L in AQP3 mutations were inserted using the QuikChange II kit (Agilent, Santa Clara, CA, USA), following the manufacturer’s instructions. For mutations L84W, L118W, L218W, L84M, L118M and L218M were introduced using the Q5 Site-Directed Mutagenesis Kit (E0554S), (New England BioLabs) according to manufacturer’s instructions.

The following primer pairs (itemised in Supplemental Table S2) were used to insert the named TbAQP2 mutations: for S131P, primers HDK1062 and HDK1063; in combination with mutation S263A, using primers HDK1064 and HDK1065 to produce plasmid pHDK166. For I110W, primers HDK607 and HDK608, to produce pHDK84; for L264R, primers HDK609 and HDK610 to produce pHDK167; the combination I110W/L264R was produced using primers HDK609 and HDK610 on plasmid pHDK84 to give plasmid pHDK78; for L258Y, primers HDK1109 and HDK1110 to produce pHDK168; for I190T, primers HDK1056 and HDK1057 to produce pHDK163; for W192G, primers HDK1058 and HDK1059 to produce pHDK164; the combination I190T/W192G was produced using primers HDK1060 and HDK1061 on plasmid pHDK163 to give plasmid pHDK165; for L84W, primers HDK1276 and HDK1277, producing pHDK210; for L118W, primers HDK1274 and HDK1275, producing pHDK208; for L218W, primers HDK1272 and HDK1273, producing pHDK209; for the combination L84W/L118W, primers HDK1276 and HDK1277 on template pHDK208, producing pHDK227; for L84M, primers HDK1364 and HDK1367, producing pHDK234; for L118M, primers HDK1365 and HDK1367, producing pHDK235; and for L218M, primers HDK1366 and HDK1367, producing pHDK236. To insert the named mutations into TbAQP3, the following primers were used (Supplemental Table S3): for W102I, primers HDK511 and HDK512, in combination with mutation R256L, with primers HDK513 and HDK514, to produce plasmid pHDK71; and to add mutation Y250L to this combination, primers HDK795 and HDK796, to produce pHDK121. All plasmids were checked by Sanger Sequencing (Source BioScience, Nottingham, UK) for the presence of the correct mutation(s) and the cassette for integration digested out with *Asc*I (NEB, Hitchin, UK) prior to transfection.

For transfection, 10 µg of digested plasmid and 1-2×10^7^ parasites of the desired cell line (either *aqp2/aqp3* null or *aqp1/aqp2/aqp3* null) were resuspended in transfection buffer and transfected using an Amaxa Nucleofector, with program X-001. After a recovery period (8-16 h) in HMI-11 at 37 °C and 5% CO_2_, the parasites were cloned out by limiting dilution with the selection antibiotic (2.5 µg/ml hygromycin). In all cases the correct integration of the expression cassettes was analysed by PCR.

### Drug sensitivity assays

Drug sensitivity assays for *T. b. brucei* bloodstream forms used the cell viability dye resazurin (Sigma) and were performed exactly as described (Wallace *et al,* 2002; Bridges *et al,* 2007) in 96-well plates with doubling dilutions of test compound, starting at 100 µM, over 2 rows of the plate (23 dilutions plus no-drug control). Incubation time with test compound was 48 h (37 °C/5% CO_2_), followed by an additional 24 h in the presence of the dye.

### Molecular dynamics

Molecular dynamics simulations were performed using the GROMACS software package, version 5.1.1 (Abraham *et al,* 2015). We used the coordinates from the homology model of TbAQP2 published in Figure 2A in Munday et al (2015a), which was inserted into POPC/POPE (4:1) membranes, approximately reflecting the membrane composition of *T. b. brucei* (Smith & Bütikofer, 2010). The membrane models were constructed using the CHARMM-GUI webserver (Jo *et al,* 2008). Subsequently, extended stability tests of the modelled structure and the bound pentamidine were carried out using unbiased simulations of 100 ns length. The root-mean-square deviation (RMSD) of the protein remained relatively low with a backbone RMSD converging to ∼3 Å after 100 ns simulated time (Supplemental Fig. S6) bound to the binding site defined previously using molecular docking (Fig. 6) (Munday *et al,* 2015a). For these and all following simulations, we used the CHARMM36 force field (Klauda *et al,* 2010); pentamidine was parameterised using the CHARMM generalized force field approach (CHGenFF (Vanommeslaeghe *et al,* 2010)). All simulations employed a time step of 2 fs under an NPT ensemble at p = 1 bar and T = 310 K. To obtain non-equilibrium work values for removing pentamidine from the internal AQP2 binding site, we then conducted steered MD simulations with a probe speed of 0.005 nm/ns and a harmonic force constant of 300 kJ/mol nm^2^, pulling pentamidine in both directions along the pore axis. The free energy profile of pentamidine binding to the AQP2 pore was reconstructed by using the Jarzynski equality (Jarzynski, 1997).

### Statistical analysis

All transport experiments were performed in triplicate and all values such as rate of uptake, percent inhibition, K_i_, K_m_, V_max_ etc were performed at least three times completely independently. For drug sensitivity tests, all EC_50_ values were based on serial dilutions over two rows of a 96-well plate (23 doubling dilutions plus no-drug control), which were obtained independently at least three times. EC_50_ and IC_50_ values were determined by non-linear regression using the equation for a sigmoid curve with variable slope and are presented as average ± SEM. Statistical significance between any two data points was determined using Student’s t-test (unpaired, two-tailed).

## Acknowledgements

This work was supported by the UK Medical Research Council (MRC) [grant 84733 to HPdK] and by the US National Institutes of Health (NIH) [Grant No. GM111749 to DWB]. DC was supported by an MRC iCASE award [MR/R015791/1]. UZ acknowledges funding from the Scottish Universities Physics Alliance. AHA is funded through a PhD studentship from Albaha University, Saudi Arabia. GDC was funded by a PhD Studentship from Science Without Borders [206385/2014-5, CNPq, Brazil]. The authors thank Dr Tansy Hammarton for the use of the CRK2 RNAi cell line and Prof David Horn for the use of the aqp1-3 null cell line.

## Author contributions

Performed experiments: AHA, JCM, GDC, MEMA, PM, LW, DP, AD, JW, GS, LFA, SPYK, HMSI, MIAS, AAE, IAT, SG, HPDK

Chemical synthesis and SAR: CEO, AK, FH, CD

Computational analysis and modelling: DG, FS, LS, CMW, MC

Supervision: RRT, DWB, PMON, UZ, HPDK

Writing manuscript: FH, UZ, HPDK

## Supplemental Materials

**Figure S1.**
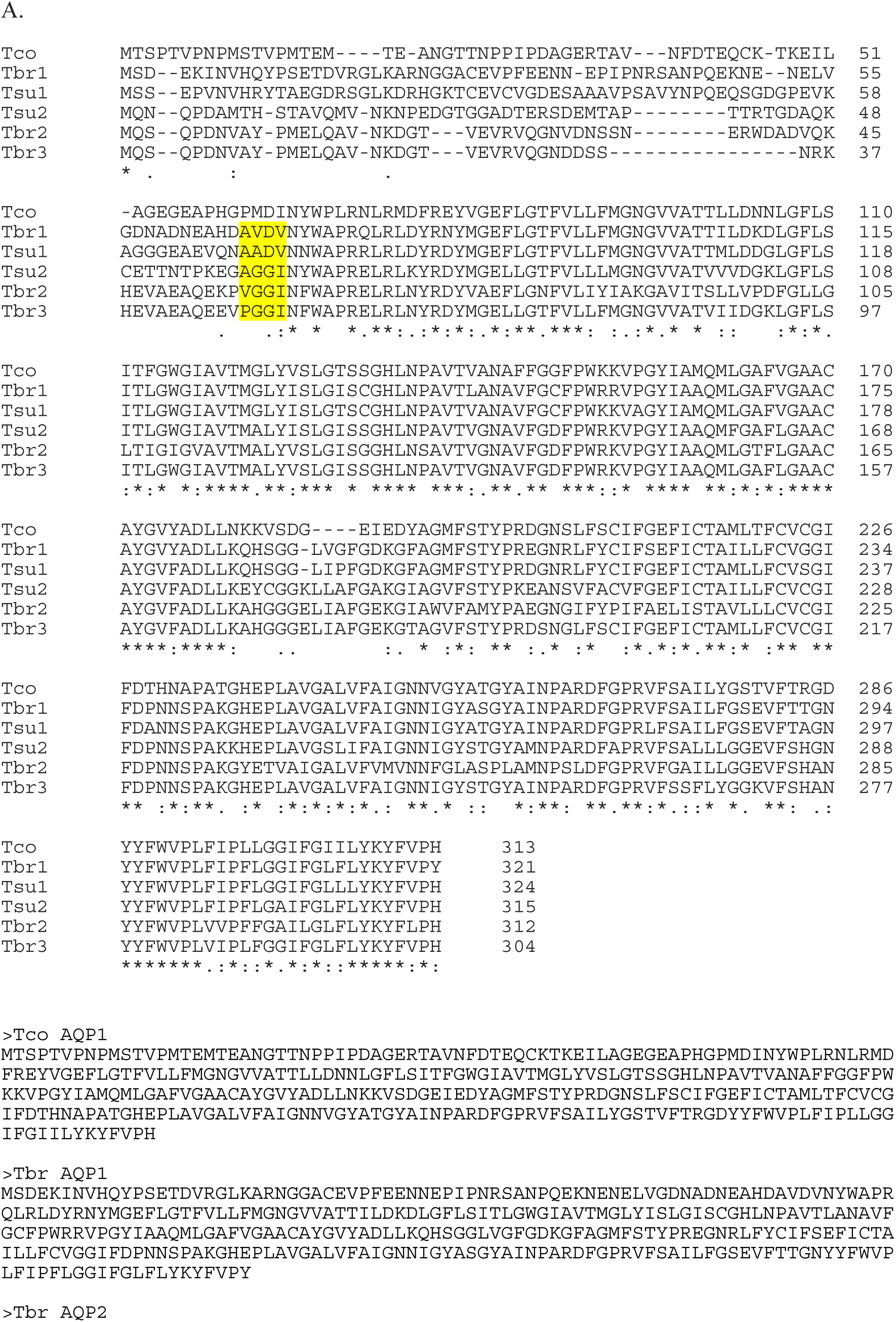
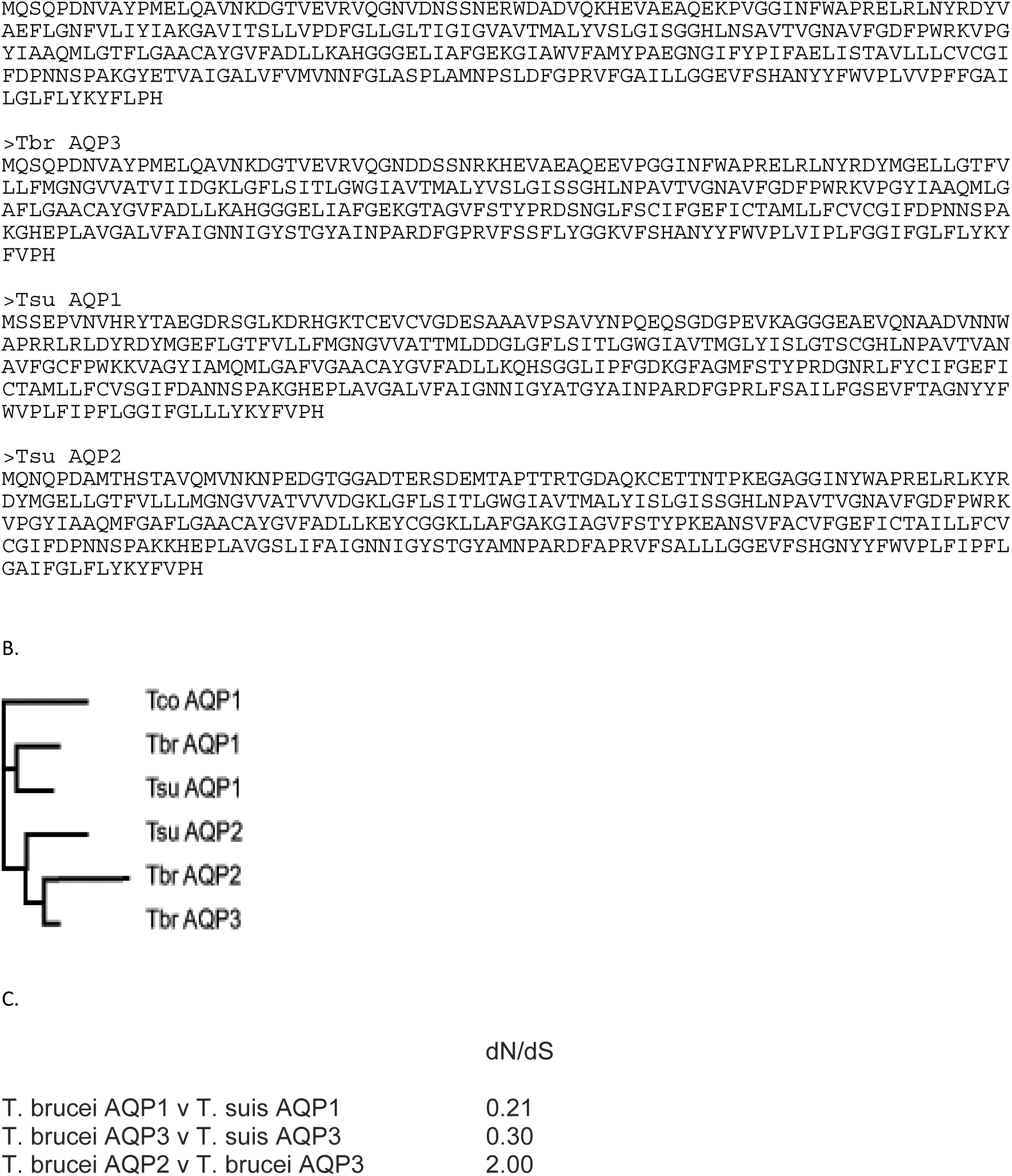
A. Sequence alignment and individual sequences of the *T. congolense*, *T. b. brucei* and *T. suis*. The *T. brucei* and *T. congolense* sequences were obtained from tritrypDB, *T. suis* sequences (Kelly S, Gibson W and Carrington M. The genome of *Trypanosoma suis*. In preparation). The alignment was produced with Clustal Omega. The yellow highlighting indicates the N-terminus of the sequences used to determine non-synonymous v synonymous ratios. B. Phylogenetic tree of these sequences. The tree is a Neighbour-joining tree produced in Clustal Omega with the lengths of the horizontals proportional to the differences. C. The ratio of non-synonymous v synonymous (dN/dS) codon changes calculated for selected comparisons between *T. brucei* and *T. suis* AQPs. The ratios were calculated using a region of high confidence alignments from ∼amino acid 60 (highlighted in Supplemental Figure 2A) to the C-terminus.

**Fig. S2.**
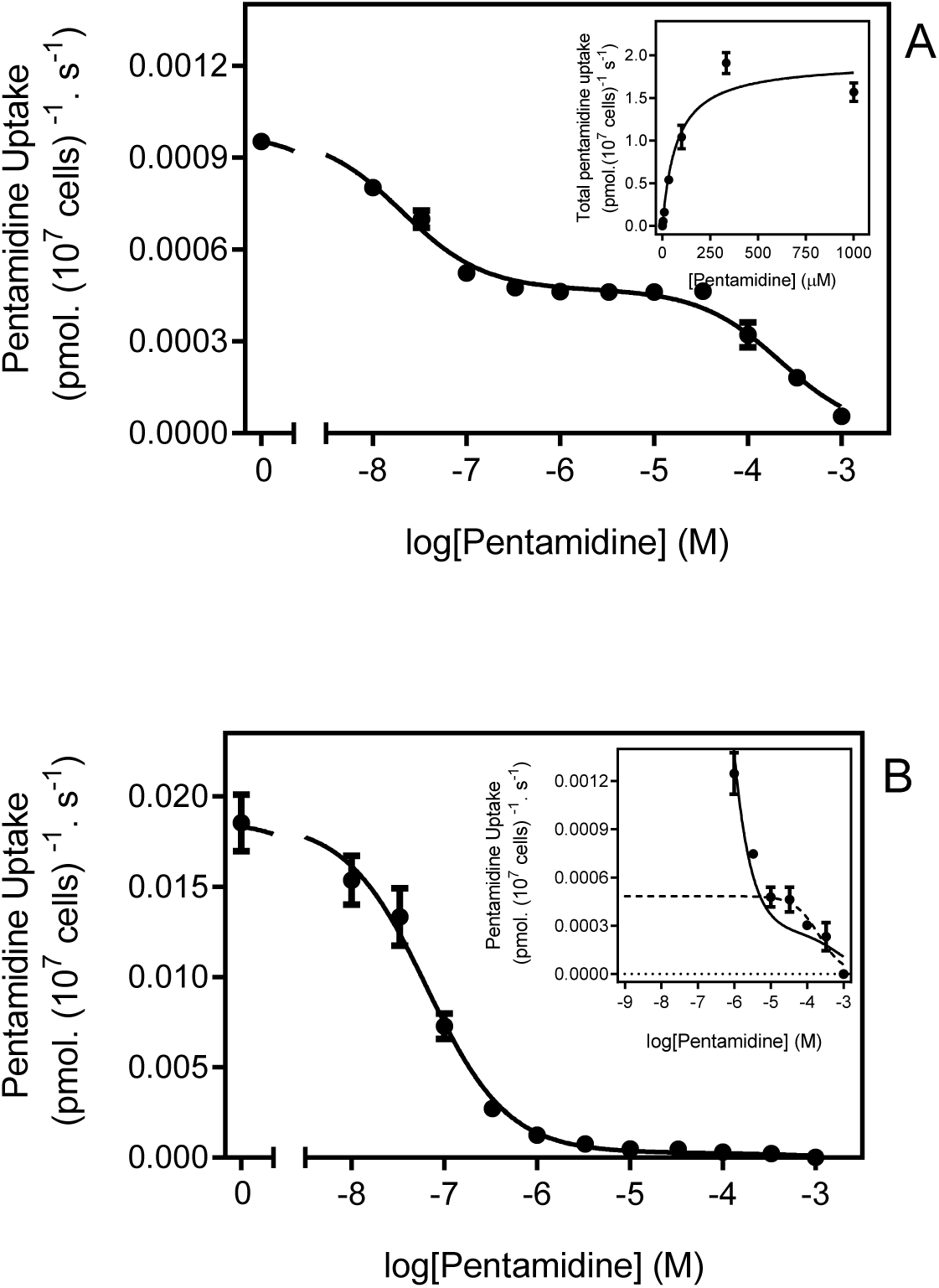
**Pentamidine transport analysis for TbAQP2^I190T^ and TbAQP2WT**. (A) Transport of 30 nM [^3^H]-pentamidine by *tbaqp2/tbaqp3* null cells expressing TbAQP2^I190T^, in the presence of unlabelled pentamidine at the indicated concentrations. Incubation time was 15 min, required to ensure sufficient radiolabel for accurate quantification, and uptake was linear and through zero over this period. The inhibition data were plotted to a double sigmoidal curve (Prism 7.0) with the bottom value fixed at 0. The high affinity component displayed an average an IC_50_ of 30.9 ± 12.2 nM (n=3) and the lower affinity segment could be converted to a Michaelis-Menten plot for determination of K_m_ and V_max_ (inset), yielding an average K_m_ of 59.9 ± 9.1 µM (n=3). The plot shown is one representative experiment in triplicate of three independent experiments. (B) Like (A) but with *tbaqp2/tbaqp3* null cells expressing TbAQP2WT. Incubation time was 20 s (linear phase). The high affinity phase had statistically identical EC_50_ (41 ± 17 nM; P>0.05) as TbAQP2^I190T^. The inset shows a zoom-in on the low-affinity part of the curve, with the dotted line representing a theoretical sigmoid plot for 1 inhibitor, with the upper limit fixed at the value obtained for 10 μM pentamidine. The low affinity component was also statistically identical in the two strains (TbAQP2-WT K_m_ = 82.7 ± 17.5 µM (n=3; P>0.05)). Note that the amount of [^3^H]-pentamidine taken up by the low affinity component is highly similar for the mutant (A) and control (B) cell lines, at approximately 0.0005 pmol(10^7^ cells)^-1^s^-1^. Both frames show one representative experiment of three repeats, each performed in triplicate. Error bars are SEM, when not shown, fall within the symbol.

**Figure S3.**
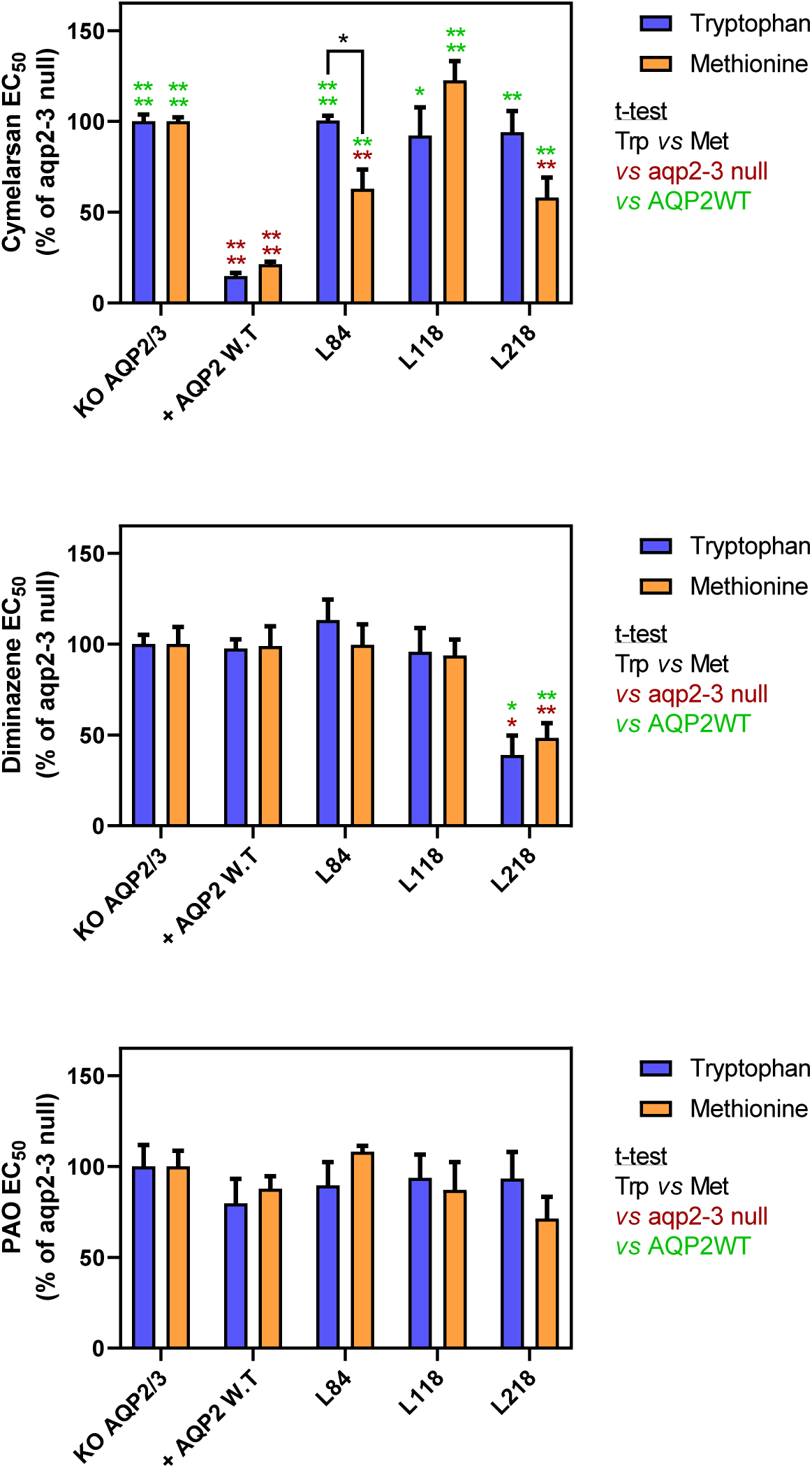
EC_50_ values for Cymelarsan, diminazene aceturate and phenylarsine oxide (PAO) against the *tbaqp2-tbaqp3* null cell line. AQP2-WT and various mutant versions thereof (indicated) were expressed in this cell line. EC50 values were determined using the alamar blue (resazurin) assay. Bars represent the average and SEM for at least three determinations. nd, not done.

**Figure S4.**
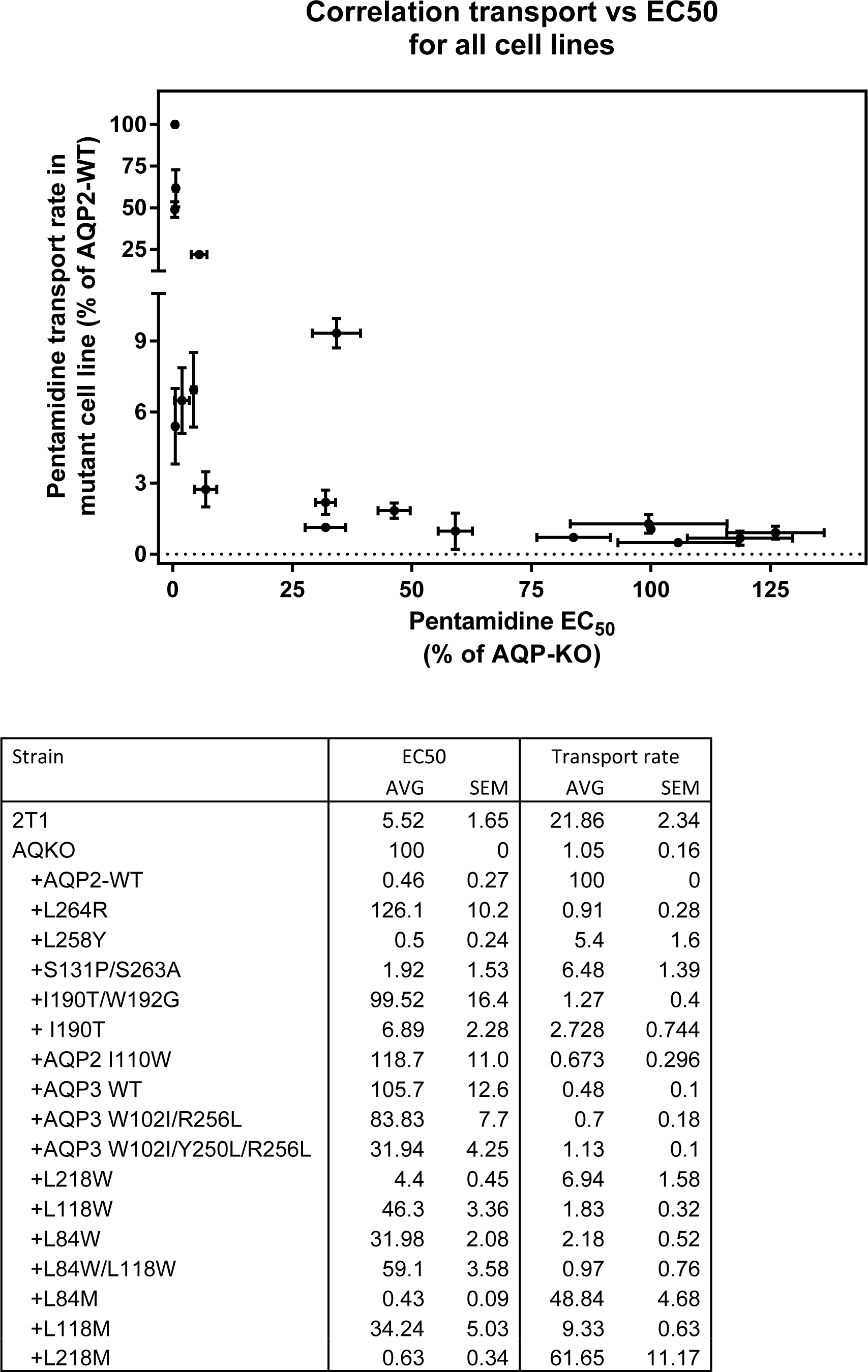
Correlation of the EC_50_ value with the rate of pentamidine transport for all 19 cell lines expressing a wild-type or mutant TbAQP2 in the aqp2/3 null *T. b. brucei line*. All EC_50_ values are expressed as percentage of the resistant control, aqp2/3 null transfected with an empty vector (no TbAQP2). 2T1 is the parental cell line of the aqp2/3 null. All values are the average of at least three independent determinations; the sensitive and resistant control cell lines were included in each independent experiment and the percentages taken are from the internal control rather than from the grand average over all experiments.

**Figure S5.**
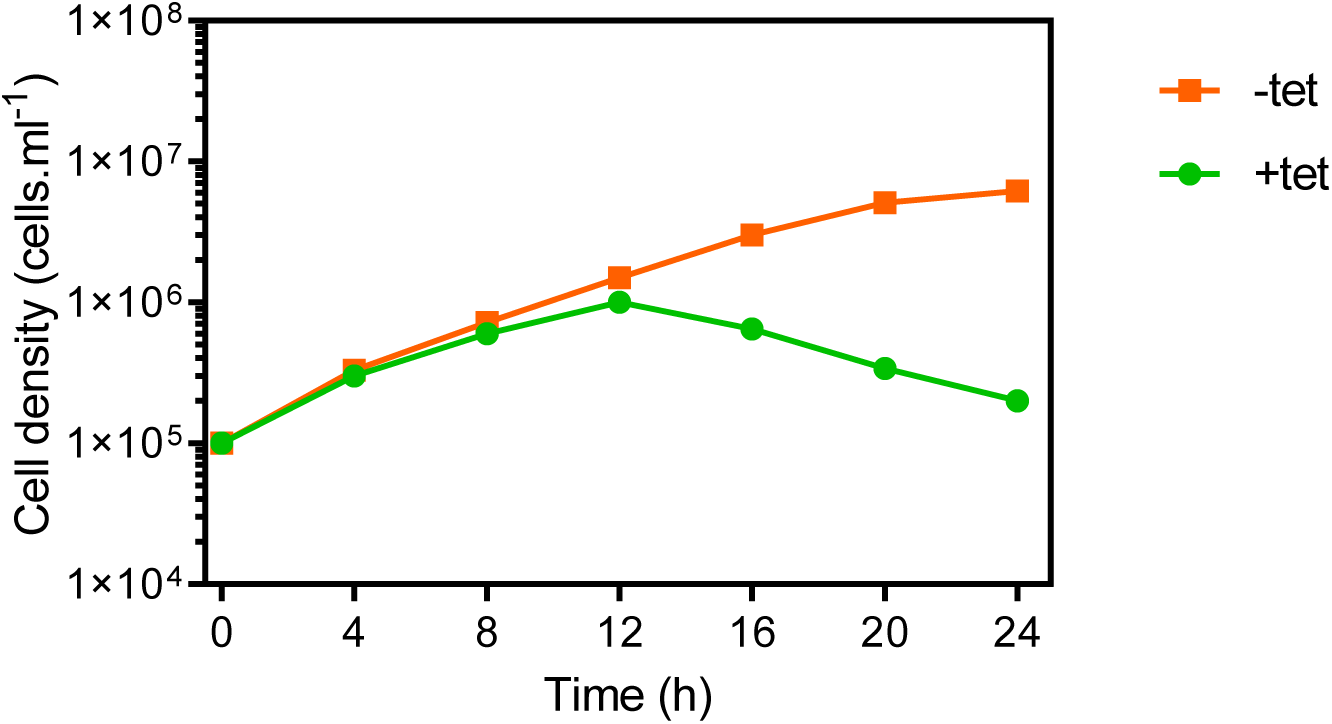
Growth Curve of CRK12 RNAi cells in full HMI-9 medium at 37 °C/5% CO_2_, in the presence or absence of 1 μg/ml tetracycline (tet). Cell counts were performed with a haemocytometer and the average of duplicate determinations is shown.

**Figure S6.**
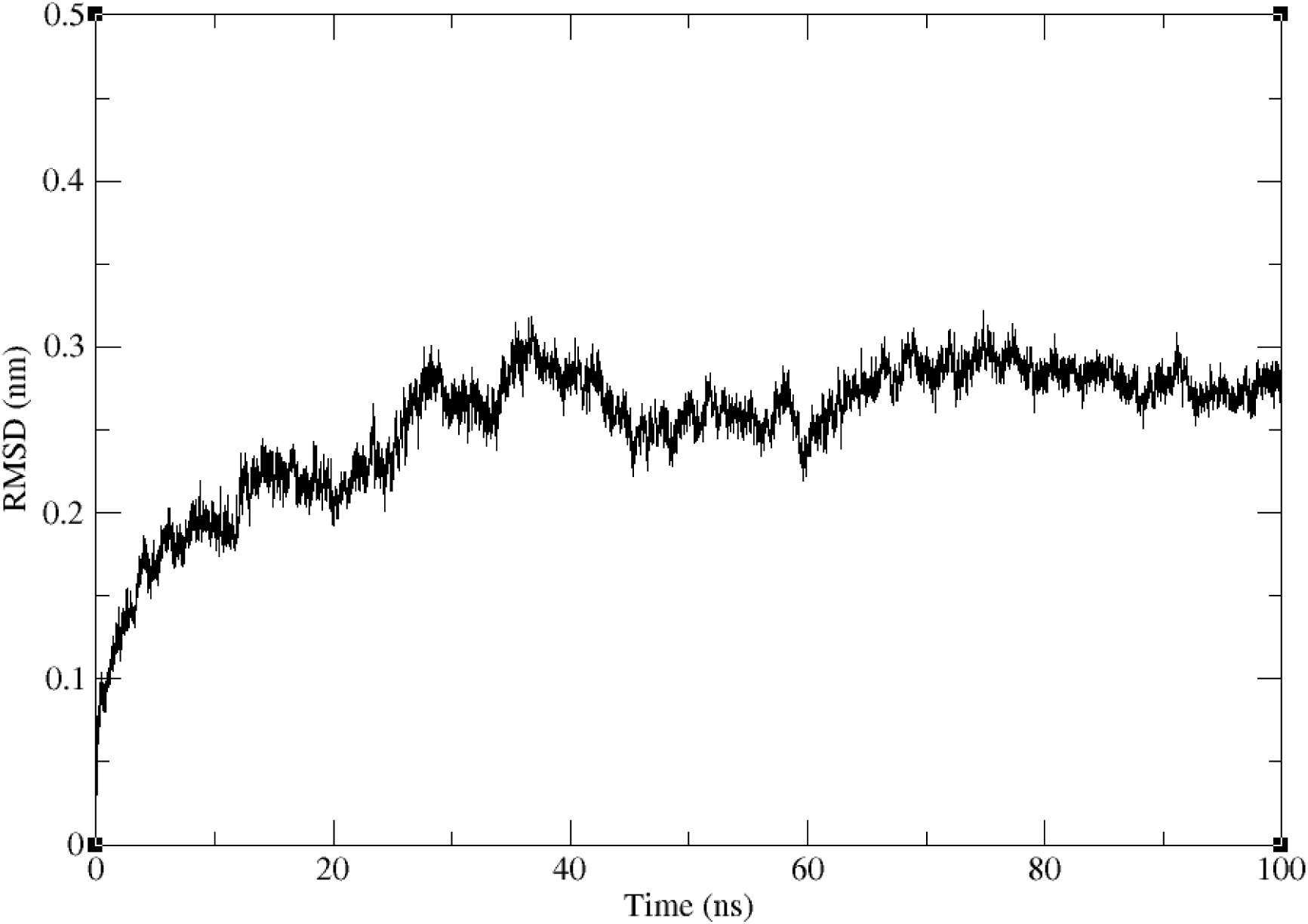
Backbone RMSD of the protein inserted into a lipid bilayer showing convergence to ∼3 Å in a simulation of 100 ns length.

**Figure S7.**
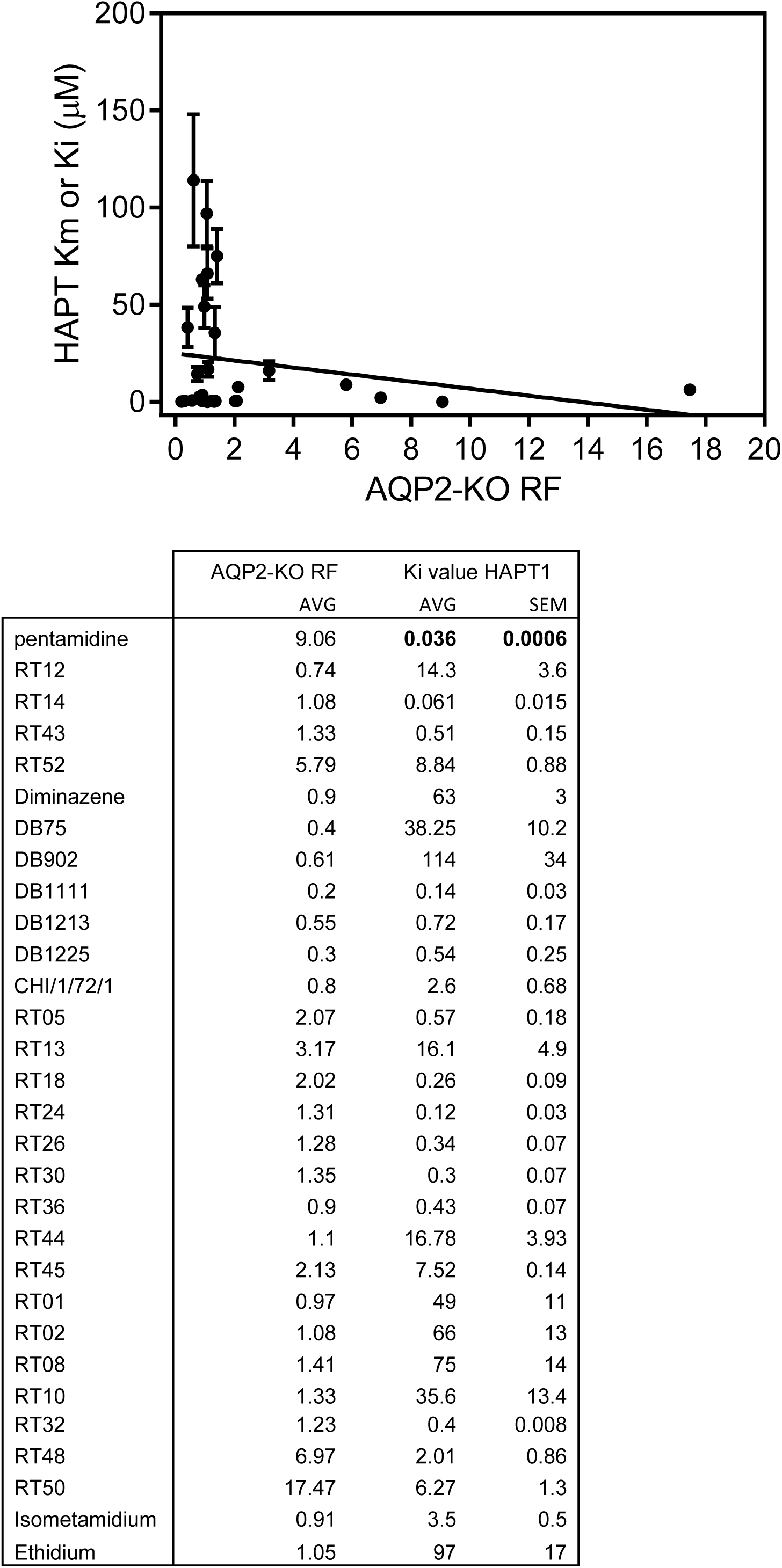
Correlation between the Resistance Factor (RF; EC_50_(aqp2/3 null)/EC_50_(TbAQP2-WT)) and the K_i_ value for inhibition of the High Affinity Pentamidine Transporter (HAPT1) encoded by TbAQP2. The pentamidine value (bold) is the K_m_ determined with radiolabeled pentamidine. The table lists the data points shown in the plot. The line was made by linear regression (Prism 6.0); correlation coefficient r^2^ is 0.039. F-test: slope is not significantly different from zero (*P* = 0.29).

**Supplemental Table S2:**
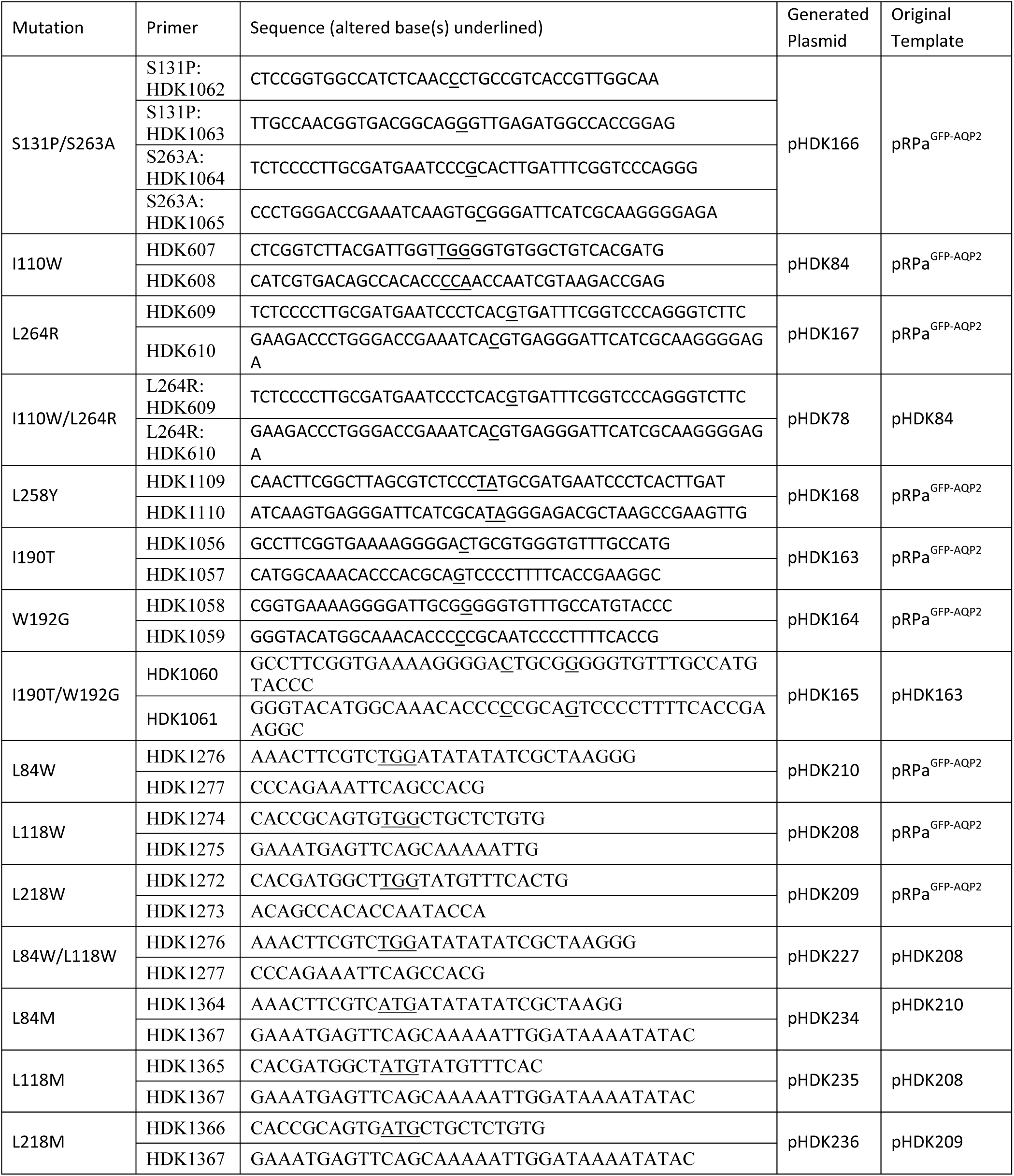
primers used for mutations in TbAQP2.

**Supplemental Table S3:**
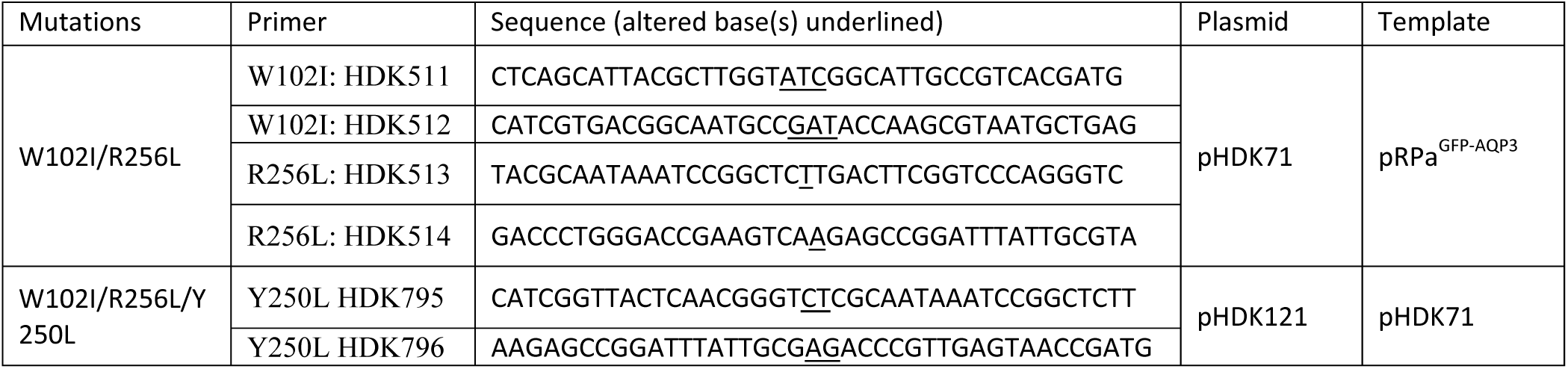
primers used for mutations in TbAQP3.

## Chemistry of new compounds from Paul O’Neill laboratory

**1,2-Bis (4-cyanophenoxy) ethane** (Ethamidine Precursor) **(1a)** ^1^

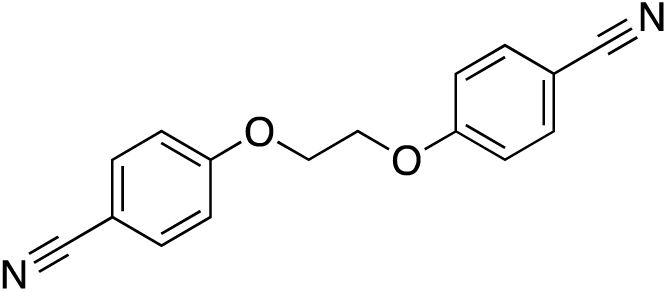

Sodium (0.16 g, 6.96 mmol) was added portionwise to anhydrous EtOH (4.0 mL) under an atmosphere of nitrogen. After dissolution of the sodium pieces, a solution of 4-cyanophenol (0.75 g, 6.38 mmol) dissolved in anhydrous EtOH (4.0 mL) was added followed by dropwise addition of 1,2-dibromoethane (0.28 mL, 3.19 mmol). The reaction mixture was allowed to stir at reflux under a nitrogen atmosphere for 3 days after which the mixture was cooled, filtered, the solid washed with water and dried under vacuum. Purification by column chromatography eluting with DCM: hexane (8:2) gave the desired dinitrile ethamidine precursor (**1a**) as a white solid (1.42 g, 84%). Mp 211-212°C; ^1^H NMR (CDCl_3_, 400MHz) δ 7.61 (d, 4H, *J* = 9.0 Hz, ArH), 7.01 (d, 4H, *J* = 9.0 Hz, ArH), 4.39 (s, 4H, CH_2_); ^13^C NMR (CDCl_3_, 100MHz) δ 161.6, 134.1, 118.9, 115.3, 104.7, 66.4; ν_max_ (NujOI) /cm^-1^ 3326 (C-O-C), 3033 (ArH), 2898 (OH), 2223 (CN), 1602 (Ar), 1509 (Ar), 1247 (C-O-C); *m/z* (CI) 282 ([M+NH_4_]^+^), found 282.12433, C_16_H_16_O_2_N_3_ requires 282.12424; anal. Found C 72.37, H 4.51, N 10.54, C_16_H_12_O_2_N_2_ requires C 72.71, H 4.57, N 10.60.

**4,4’-(ethane-1,2-diylbis(oxy))dibenzimidamide dihydrochloride dihydrate** (Ethamidine, Compound CHI/1/30/1) (**2**) ^1^

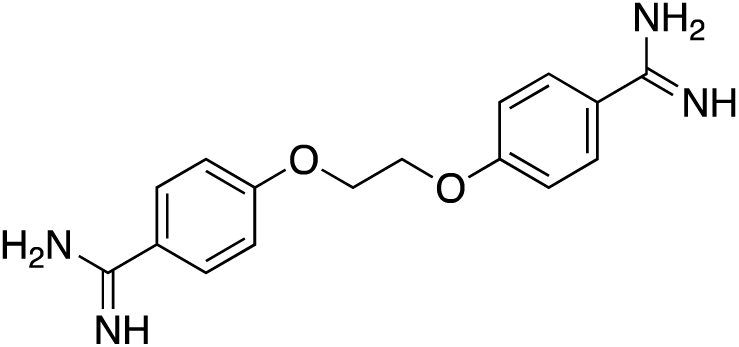

(0.51 g, 1.92 mmol) of **1a** was dissolved in a mixture of anhydrous benzene (54 mL) and EtOH (2.90 mL), cooled to 0 °C and saturated with HCl gas. The mixture was sealed and allowed to stir at room temperature for 3 days after which anhydrous Et_2_O (28 mL) was introduced and the mixture was allowed to stir for 10 minutes. The solids were filtered under nitrogen and dissolved in a mixture of anhydrous EtOH (36 mL) and EtOH.NH_3_ (36 mL). The mixture was heated overnight (50 °C), cooled to room temperature and reduced by half *in vacuo*. Ether was added to precipitate the solid which was filtered, washed and dried under vacuum. Purification by recrystallisation (2N HCl) gave the desired compound **2** as fine white needles (0.54 g, 70%). Mp 333°C; ^1^H NMR (MeOD, 400MHz) δ 7.85 (d, 4H, *J* = 9.0 Hz, ArH), 7.23 (d, 4H, *J* = 9.0 Hz, ArH), 4.52 (s, 4H, CH_2_); ^13^C NMR (MeOD, 100MHz) δ 167.9, 165.3, 131.5, 121.7, 116.7, 68.5; ν_max_ (Nujol) /cm^-1^ 3362 (NH), 3037 (ArH), 2940 (C-H), 1658 (C=N-H), 1606 (Ar), 1505 (Ar), 1245 (C-0-C); *m/z* (ESP) 299 ([M-H]^-^); anal. Found C 47.54 H 5.62 N 14.30, C_16_H_24_N_4_O_4_Cl_2_ requires C 47.18, H 5.94, N 13.76.

**4, 4’-(propane-1,3-diylbis(oxy))dibenzonitrile** (Propamidine Precursor) (**1b**)^1^

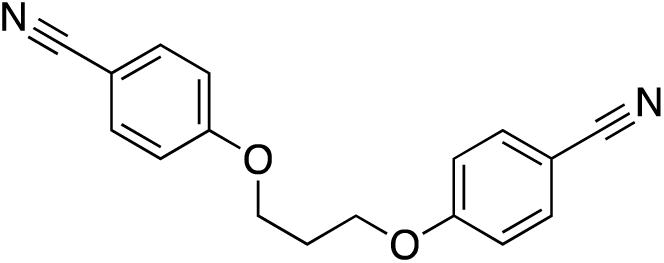

Sodium (0.16 g, 6.96 mmol) was added portionwise to anhydrous EtOH (4.0 mL) with stirring under an atmosphere of nitrogen. After dissolution of Na, a solution of 4-cyanophenol (0.75 g, 6.38 mmol) dissolved in dry ethanol (4.0 mL) was added followed by dropwise addition of 1,3-dibromopropane (0.32 mL, 3.19 mmol). The reaction mixture was allowed to stir at reflux under a nitrogen atmosphere for 3 days after which the mixture was cooled, filtered, the solid washed with water and dried under vacuum. Purification by column chromatography eluting with DCM: hexane (8:2) gave the desired compound **1b** as a white solid (1.40 g, 79%). Mp 190-191°C; ^1^H NMR (CDCl_3_, 400MHz) δ 7.59 (d, 4H, *J* = 8.5 Hz, ArH), 6.96 (d, 4H, *J* = 8.5 Hz, ArH), 4.20 (t, 4H, *J* = 6.0 Hz, CH_2_), 2.32 (m, 2H, CH_2_); ^13^C NMR (CDCl_3_, 100MHz) δ 161.9, 134.0, 119.1, 115.1, 104.2, 64.4, 28.8; ν_max_ (Nujol) /cm^-1^ 3104 (Ar-H), 2823 (C-H), 2221 (C≡N), 1604 (Ar), 1509 (Ar), 1253 (C-O); *m/z* (CI) 296 ([M+NH_4_]^+^), found 296.14037, C_17_H_18_N_3_O_2_ requires 296.13992; anal. Found C 73.22, H 5.13, N 10.03, C_17_H_14_N_2_O_2_ requires C 73.37, H 5.07, N 10.07.

**4,4’-(propane-1,3-diyl*bis*(oxy))dibenzimidamide dihydrochloride dihydrate** (Propamidine, Compound CHI/1/25/5) (**3**)^1^

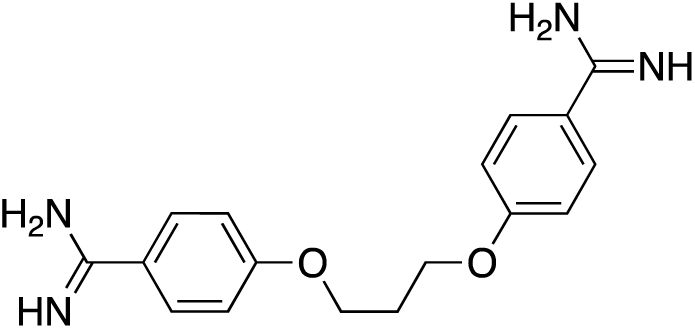

(0.50 g, 1.79 mmol) of **1b** was dissolved in a mixture of anhydrous benzene (55 mL) and anhydrous ethanol (3.0 mL), cooled to 0 °C and saturated with HCl gas. The mixture was sealed and allowed to stir at room temperature for 3 days after which ether (30 mL) was added and the mixture was allowed to stir for 10 minutes. The solids were filtered under nitrogen and dissolved in a mixture of anhydrous EtOH (36 ml) and EtOH.NH_3_ (36 mL). The mixture was heated overnight (50 °C), cooled to room temperature and reduced by half *in vacuo*. Et_2_O (15 mL) was added to precipitate the solid which was filtered, washed and dried under vacuum. Purification by recrystallisation (2N HCl) gave the desired compound **3** as fine white needles (0.59 g, 78%). Mp 200°C; ^1^H NMR (MeOD, 400MHz) δ 7.82 (d, 4H, *J* = 9.0 Hz, ArH), 7.19 (d, 4H, *J* = 9.0 Hz, ArH), 4.34 (t, 4H, *J* = 6.0 Hz, CH_2_), 2.46 (m, 2H, CH_2_); ^13^C NMR (MeOD, 100MHz) δ 167.9, 165.5, 131.5, 121.5, 121.4, 116.6, 66.4, 30.3; ν_max_ (Nujol) /cm^-1^ 3280 (N-H), 3038 (Ar-H), 2929 (C-H), 1504 (Ar), 1606 (Ar), 1240 (C-O-C); *m/z* (ESP) 313 ([M-H]^-^); anal. Found C 48.60, H 6.10, N 13.25, C_17_H_26_N_4_O_4_Cl_2_ requires C 48.46, H 6.22, N 13.30.

**4,4’-(Butane-1,4-diylbis(oxy))dibenzonitrile** (Butamidine precursor) (**1c**)^1^

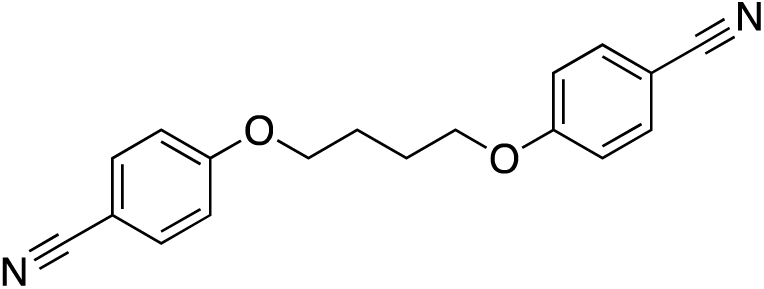

Sodium (0.10 g, 4.35 mmol) was added portionwise to dry EtOH (4.0 mL) stirring under an atmosphere of nitrogen. After dissolution of sodium, a solution of 4-cyanophenol (0.47 g, 3.95 mmol) dissolved in dry ethanol (4.0 mL) was added followed by dropwise addition of 1,4-dibromobutane (0.24 mL, 1.98 mmol). The reaction mixture was allowed to stir at reflux under a nitrogen atmosphere for 3 days after which the mixture was cooled, filtered, the solid washed with water and dried under vacuum. Purification by column chromatography eluting with DCM: hexane (8:2) gave the desired compound **1c** as a white solid (1.03 g, 89%). Mp 174°C; ’H NMR (CDCI_3_, 400MHz) δ 7.59 (d, 4H, *J* = 8.9 Hz, ArH), 6.93 (d, 4H, *J* = 8.9 Hz, ArH), 4.08 (m, 4H, CH_2_), 2.01 (m, 4H, CH_2_); ^13^C NMR (CDCl_3_, 100MHz) δ 162,1, 134.0, 119.1, 115.1, 104.0, 67.7, 25.7; ν_max_ (Nujol) /cm^-1^ 3332 (C-O-C), 3033 (Ar-H). 2956 (C-H), 2219 (C≡N), 1604 (Ar). 1506 (Ar), 1251 (C-O-C); *m/z* (CI) 310 ([M+NH_4_]^+^) found 310.15532, C_18_H_20_N_3_O_2_ requires 310.15555; anal. Found C 74.03, H 5.55, N 9.55, C_18_H_16_N_2_O_2_ requires 73.95, H 5.52, N 9.58.

**4,4’-(Butane-l,4-diylbis(oxy))dibenzimiamide dihydrochloride dihydrate** (Butamidine, Compound CHI/1/41/1) (**4**)^1^

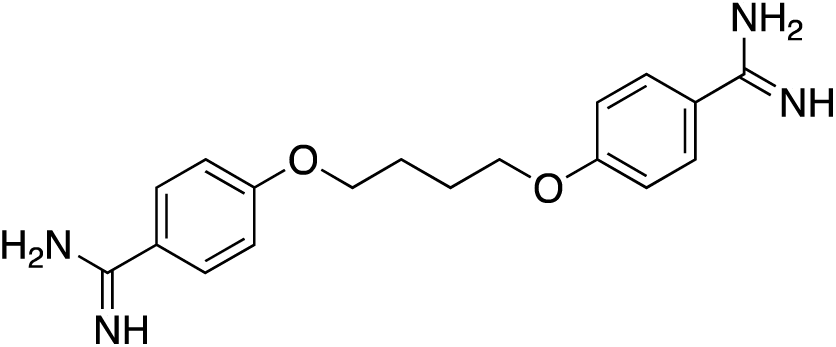

Compound **1c** (0.42 g, 1.44 mmol) was dissolved in a mixture of anhydrous benzene (46 mL) and anhydrous ethanol (2.50 mL), cooled to 0 °C and saturated with HCl gas. The mixture was sealed and allowed to stir at room temperature for 3 days after which anhydrous Et_2_O (40 mL) was introduced and the mixture was allowed to stir for 10 minutes. The solids were filtered under nitrogen and dissolved in a mixture of anhydrous EtOH (34 mL) and EtOH.NH_3_ (34 mL). The mixture was heated overnight (50 °C), cooled to room temperature and reduced by half *in vacuo*. Ether (15 mL) was added to precipitate the solid which was filtered, washed and dried under vacuum. Purification by recrystallisation (2N HCl) gave the desired compound **4** as fine white needles (0.46 g, 73%). Mp 286-287°C’, ‘H NMR (MeOD, 400MHz) δ, 7.80 (d, 4H, *J* = 9.0 Hz, ArH), 7.14 (d, 4H, *J* = 9.0Hz, ArH), 4.19 (m, 4H, CH_2_), 2.02 (m, 4H, CH_2_); ^13^ CNMR (MeOD, 100MHz) δ 165.7, 131.4, 121.2, 116.7, 69.7, 27.2; ν_max_ (Nujol) /cm^-1^ 3370 (N-H), 3129 (Ar-H), 2884 (C-H), 1650 (C≡N), 1606 (Ar), 1508 (Ar), 1257 (C-O); *m/z* (ESP) 327 ([M+H]^+^); anal. Found C 49.87, H 6.46, N 12.67, C_18_H_28_N_4_O_4_Cl_2_ requires C 49.66, H 6.48, N 12.87.

**4-(4-phenoxybutoxy) benzonitrile**

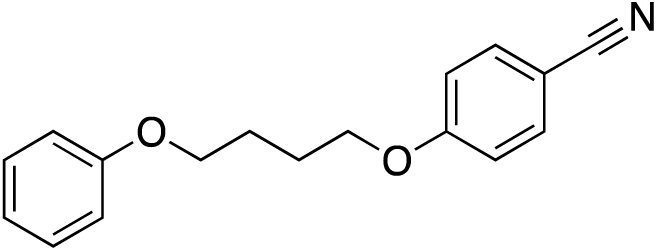

Sodium (0.16g, 6.96 mmol) was added dropwise to dry ethanol (5 mL) and dissolved under a nitrogen atmosphere. To this a solution of 4-cyanophenol (0.53g, 4.47 mmol) dissolved in anhydrous ethanol (5ml) was added followed by addition of 1,4-dibromobutane (0.53 mL, 4.47 mmol). The reaction mixture was allowed to stir at reflux and monitored by TLC. After consumption of 4-cyanophenol, the reaction mixture was allowed to cool to room temperature. In a separate flask sodium (0 16 g, 6.96 mmol) was added portionwise to ethanol (5 mL) stirring under nitrogen. A solution of phenol (0.42 g, 4.47 mmol) in ethanol (5ml) was added and stirred for 10 minutes. This mixture was added dropwise to the cooled mixture and allowed to stir under reflux for 3 days after which the mixture was cooled, filtered, the solid washed with water and dried under vacuum. Purification by column chromatography eluting with DCM: hexane (8:2) gave the desired compound 4-(4-phenoxybutoxy) benzonitrile as a white solid (0.98 g, 82%). Mp 130°C; ^1^H NMR (CDCl_3,_ 400MHZ) δ 7.57 (d, 2H, *J* = 8.9 Hz, ArH), 7.28 (d, 1H, *J* = 7.5 Hz, ArH), 7.26 (d, 1H, *J* = 8.1 Hz, ArH), 6.92 (m, 5H, ArH), 4.08 (t, 2H, *J* = 5.9 Hz, CH_2_), 4.03 (t, 2H, *J* = 5.9 Hz, CH_2_), 1.99 (m, 4H, CH_2_); ^13^C NMR (CDCl_3_, 100MHz) δ 162.6, 159.2, 134.3, 129.8, 121.1, 115.5, 114.8, 104.3, 68.3, 67.5, 26.2; ν_max_ (Nujol) /cm^-1^ 3043 (Ar-H), 2884 (C-H), 2219 (C≡N), 1602 (Ar), 1504 (Ar), 1247 (C-O); *m/z* (CI) 285 ([M+NH_4_]^+^), found 285.16020, C_17_H_21_N_2_O_2_ requires 285.16031; anal. Found C 76.40, H 6.46, N 5.44, C_17_H_17_NO_2_ requires C 76.38, H 6.40, N 5.24.

**4-(4-Phenoxybutoxy)benzimidamide hydrochloride hydrate** (Compound CHI/1/69/1)

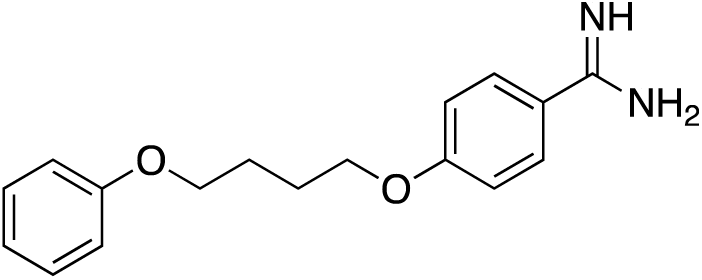

4-(4-phenoxybutoxy) benzonitrile (0.27 g, 1.01 mmol) was dissolved in a mixture of anhydrous benzene (100 mL) and ethanol (1.60 mL), cooled to 0 °C and saturated with HCl gas. The mixture was sealed and allowed to stir at room temperature for 3 days after which anhydrous Et_2_O (16 mL) was introduced and the mixture was allowed to stir for an additional 10 minutes. The solids were filtered under nitrogen and dissolved in a mixture of anhydrous EtOH (20 mL) and anhydrous EtOH.NH_3_ (20 mL). The mixture was heated overnight (50 °C), cooled to room temperature and reduced by half *in vacuo*. Ether (30 mL) was added to precipitate the solid which was filtered, washed and dried under vacuum. Purification by recrystallisation (2N HCl) gave the desired compound 4-(4**-**Phenoxybutoxy)benzimidamide hydrochloride hydrate as fine white needles (0.28 g, 82%). Mp 134-135°C; ^1^H NMR (DMSO, 400MHz) δ 9.28 (s, 2H, NH;), 9.08 (s, 2H, NHZ), 7.86 (d, 2H, *J* = 9.0 Hz, ArH), 7.29 (d, 1H, *J* = 7.0 HZ, ArH), 7.27 (d, 1H, *J* = 7.2 Hz, ArH), 7.16 (d, 2H, *J* = 9.0 Hz, ArH), 6.93 (d, 3H, *J* = 7.8 Hz, ArH), 4.16 (t, 2H, *J* = 5.9 Hz, CH_2_), 4.03 (t, 2H, *J* = 5.9 Hz, CH_2_), 1.89 (m, 4H, CH_2_); ^13^C NMR (DMSO, 100MHZ) δ 165.0, 163.3, 158.9, 130.5, 129.8, 120.7, 119.6, 115.1, 114.7, 68.1, 67.2, 25.6, 25.5; ν_max_ (Nujol) /cm^-1^ 3288 (N-H), 1656 (C=N-H), 1604 (Ar), 1506 (Ar), 1234 (C-O-C); *m/z* (ESP) 285 ([M+H]^+^), found 285.1603, C_17_H_24_N_2_O_2_ requires 285.1599; anal. Found C 59.50, H 6.75, N 8.33, C_17_H_23_N_2_O_3_Cl requires C 60.26, H 6.84, N 8.27.

**4-(5-(*p*-Tolyloxy)pentyloxy)benzonitrile**

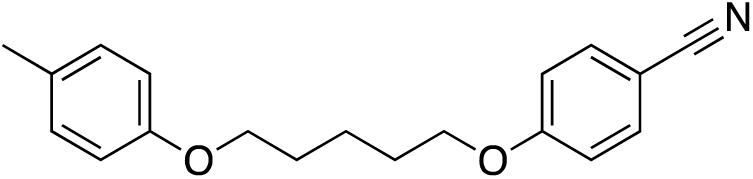

Sodium (0.12 g, 5.22 mmol) was added portionwise to anhydrous ethanol (4.0 mL) and dissolved under a nitrogen atmosphere. To this a solution of 4-cyanophenol (0.57 g, 4.79 mmol) dissolved in anhydrous ethanol (4.0 mL) was added followed by dropwise addition of 1,5-dibromopentane (0.65 mL, 4.79 mmol). The reaction mixture was allowed to stir at reflux and monitored by TLC. After consumption of 4-cyanophenol the reaction mixture was cooled to room temperature. In a separate flask, sodium (0.57 g, 4,79 mmol) was added portionwise to anhydrous EtOH (4.0 ml) with stirring under nitrogen. To this, a solution of *p*-cresol (0.5 ml, 4.79 mmol) in anhydrous EtOH (4.0 ml) was added and stirred for 10 minutes. This mixture was added dropwise to the cooled mixture and stirred under reflux for 3 days after which the mixture was cooled, filtered and the solid washed with water and dried under vacuum. Purification by column chromatography eluting with DCM: hexane (8:2) gave the desired compound as a white solid(1.02 g, 72%). Mp 133°C; ^1^H NMR (CDCl_3_, 400MHz) δ 7.57 (d, 2H, *J* = 9 Hz, ArH), 7.07 (d, 2H, *J* = 8.6 Hz, ArH), 6.93 (d, 2H, *J =* 9.0 Hz, ArH), 6.79 (d, 2H, *J* = 8.6 Hz, ArH) 4.02 (t, 2H, *J* = 6.4 Hz, CH_2_) 3.96 (t, 2H, *J* = 6.4 Hz, CH_2_), 2.28 (s, 3H, CH3), 1.86 (m, 4H, CH_2_), 1.64 (m, 2H, CH_2_); ^13^C NMR (CDCl_3_, 100MHz) δ 162.7, 157.2, 134.3, 130.3, 130.2, 115.5, 114.7, 104.1, 68.5, 68.0, 29.4, 29.1, 23.0, 20.8; ν_max_ (Nujol) /cm^-1^ 3322 (C-O-C), 3031 (Ar-H), 2921 (C-H), 2223 (C≡N), 1602 (Ar), 1506 (Ar), 1234 (C-O-C); *m/z* (CI) 313 ([M+NH_4_]^+^), found 313.19092, C_19_H_25_N_2_O_2_ requires 313.19162; anal. Found C 77.19, H 7.13, N 5.02, C_19_H_21_NO_2_ requires C 77.26, H 7.17, N 4.74.

**4-(5-(*p*-tolyloxy)pentyyloxy)benzimidamide hydrochloride hydrate** (Compound CHI/1/72/1)

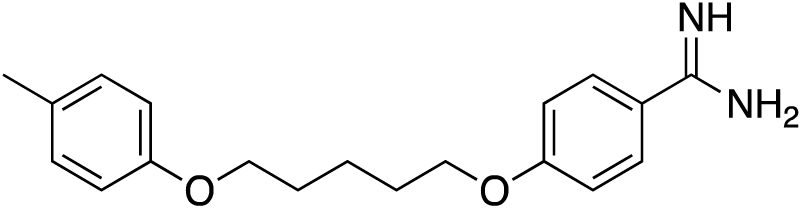

4-(5-(*p*-Tolyloxy)pentyloxy)benzonitrile (0.27 g, 0.91 mmol) was dissolved in a mixture of anhydrous benzene (100 mL) and anhydrous ethanol (1.60 mL), cooled to 0 °C and saturated with HCl gas. The mixture was sealed and allowed to stir at room temperature for 3 days after which anhydrous Et_2_O (20 mL) was introduced and the mixture was allowed to stir for 10 minutes. The solids were filtered under nitrogen and dissolved in a mixture of anhydrous EtOH (20 mL) and EtOH.NH_3_ (20 mL). The mixture was heated overnight (50 °C), cooled to room temperature and reduced by half *in vacuo*. Ether (30 mL) was added to precipitate the solid which was filtered, washed and dried under vacuum. Purification by recrystallisation (2N HCl) gave the desired compound as fine white needles (0.25 g, 75%). Mp 132°C; ^1^H NMR (DMSO, 400MHZ) δ 7.84 (d, 2H, *J* = 9.1 Hz, ArH), 7.15 (d, 2H, *J* = 9.1 Hz, ArH), 7.06 (d, 2H, *J* = 8.4 Hz, ArH), 6.80 (d, 2H, *J* = 8.4 Hz, ArH), 4.11 (t, 2H, *J* = 6.4 Hz, CH_2_), 3.93 (t, 2H, *J* = 6.4 Hz, CH_2_), 2.22 (s, 3H, CH_3_), 1.78 (m, 4H, CH_2_), 1.56 (m, 2H, CH_2_); ^13^C NMR (DMSO, 100MHz) δ 165.0, 163.3, 158.9, 130.5, 130.1, 115.1, 114.5, 79.5, 79.3, 79.0, 20.4; ν_max_ (Nujol) /cm^-1^ 3430 (NH), 3309 (C-O-C), 3093 (Ar-H), 1658 (C=N-H), 1606 (Ar), 1508 (Ar), 1245 (C-O-C); *m/z* (ESP) 313 ([M+H]^+^), found 313.1916, C_19_H_25_N_2_O_2_ requires 313.1907; anal. Found C 62.19, H 7.42, N 7.66, C_19_H_27_ClN_2_O_3_ requires C 62.20, H 7.42, N 7.66.

### Synthesis of ER 1004

**4-bromobenzothioamide**

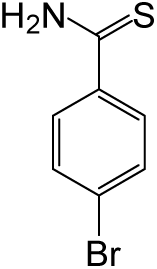

Triethylamine (3.8 mL) was added to a solution of 4-bromobenzonitrile (5 g, 27.5 mmol) in pyridine (17 mL). The solution was cooled to 10 °C and H_2_S (g) was bubbled through for 15 min. The resulting green solution was allowed to stir overnight (17 h). Nitrogen was bubbled through for 1 h to remove any excess H_2_S. Water (27 mL) was added and the mixture was stirred for 10 min, a further portion of water (62 mL) was added and the pale yellow suspension left stirring overnight. The precipitate was filtered and rinsed with water to afford the title compound as bright yellow crystals (5.52 g, 93%). ^1^H NMR (d_6_-acetone, 400MHz) 9.07 (bs, 1H, NH), 8.92 (s, 1H, NH), 7.94 (dd, 2H, *J* = 2.0, 6.5 Hz, ArH), 7.62 (dd, 2H, *J* = 2.0, 6.5 Hz, ArH); ^13^C NMR (d_6_-acetone, 100MHz) 201.9, 140.2, 132.3, 130.4, 126.5; *m/z* (CI) 216 (100%, [M^+^])

**2,4-bis(4-bromophenyl)thiazole**

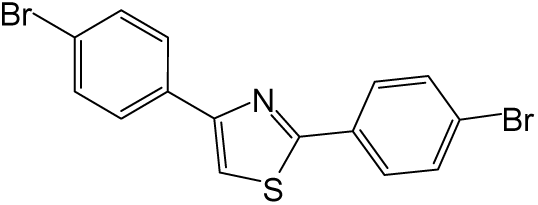

2,4’-dibromoacetophenone (1 g, 3.60 mmol) was added to a solution of 4-bromobenzothioamide (777 mg, 3.60 mmol) in EtOH (15 mL) and warmed to 45 °C for 1 h. The mixture was cooled to room temperature and left for 30 min before filtering. The precipitate was washed with EtOH: water (3:1, 10 mL) and dried to afford the thiazole as a pale solid (1.33 g, 94%). ^1^H NMR (CDCl_3_, 250MHz) 7.89 (d, 2H, *J* = 8.5 Hz, ArH), 7.85 (d, 2H, *J* = 8.5 Hz, ArH), 7.59 (d, 2H, *J* = 5.5 Hz, ArH), 7.56 (d, 2H, *J* = 5.5 Hz, ArH), 7.47 (s, 1H, CH); *m/z* (CI) 396 (10%, [M+H]^+^).

**4,4’-(thiazole-2,4-diyl)dibenzonitrile**

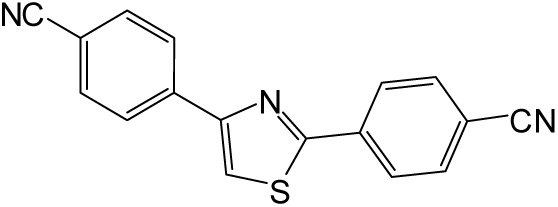

A suspension of 2,4-bis(4-bromophenyl)thiazole (1 g, 2.53 mmol) and CuCN (906 mg, 10.12 mmol) in anhydrous DMF (15 mL) were heated to reflux for 21 h. On cooling, the reaction mixture was poured into aqueous NH_4_OH (10%, 50 mL) and extracted with CHCl_3_ (100 mL). Both layers were filtered to remove the dark precipitate. The organic layer was washed with water (2 x 50 mL), brine (50 mL) and dried MgSO_4_. Removal of solvent gave a dark oily solid. Purification by column chromatography eluting with CHCl_3,_ afforded the title compound as a pale solid (361 mg, 50%). ^1^H NMR (CDCl_3_, 400MHz) 8.15 (d, 2H, *J* = 8.5 Hz, ArH), 8.11 (m, 2H, ArH), 7.77 (d, 2H, *J* = 8.5 Hz, ArH), 7.73 (t, 2H, *J* = 3.0 Hz, ArH), 7.26 (s, 1H, CH); ^13^C NMR (CDCl_3_, 100MHz) 166.5, 137.4, 133.2, 130.1, 127.3, 119.1, 117.1, 114.2, 112.4; *m/z* (CI) 288 (100 %, [M+H]^+^).

**4,4’-(thiazole-2,4-diyl)dibenzimidamide (ER1004)**

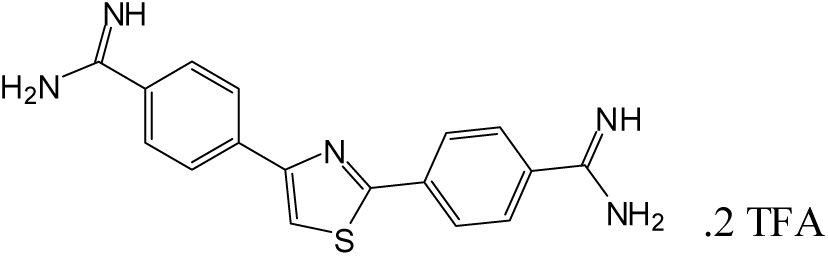

The Garigipati Reaction is a little known reaction which effects the conversion of hindered nitriles to unsubstituted amidines in a mild and effective manner ^2, 3^. This is an efficient one step transformation involving direct nucleophilic addition of an amine to a nitrile, affording the corresponding amidine (Scheme 1).

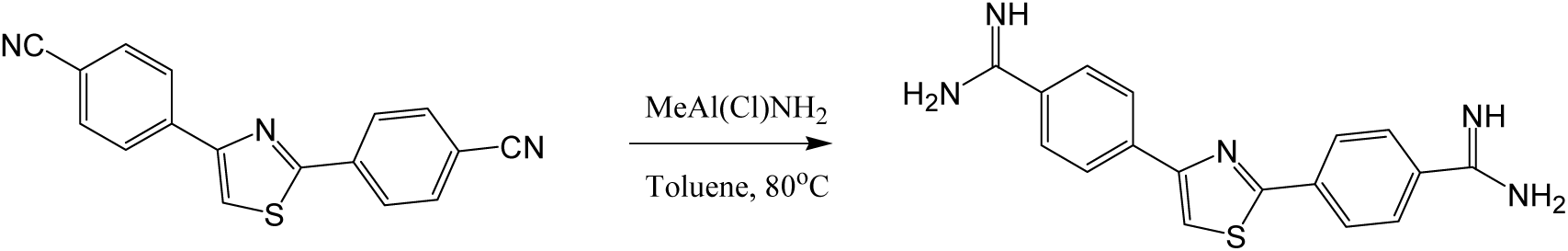

### Scheme 1

The alkylchloroaluminium amides are effectively generated from trimethyl aluminium and ammonium chloride and the intermediate aluminium complex is easily hydrolyzed by water adsorbed on silica gel (Scheme 2).

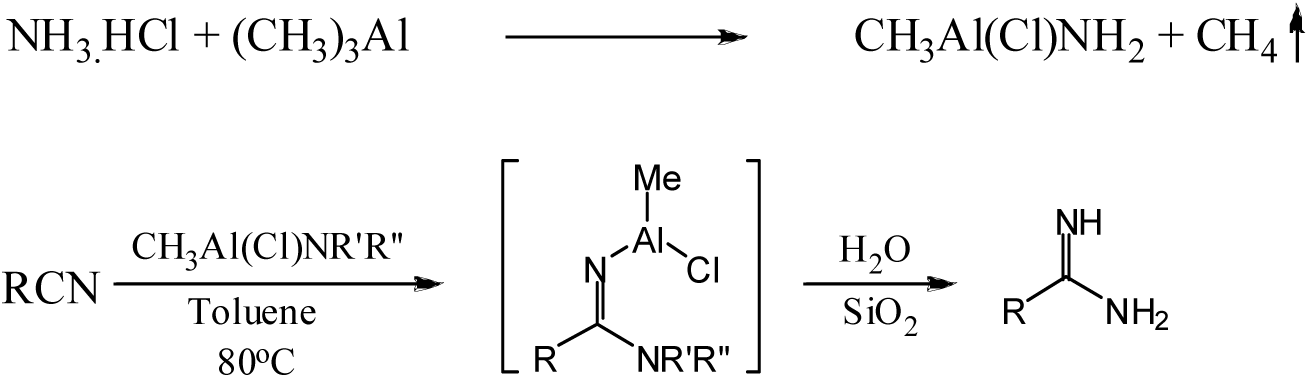

### Scheme 2

The freshly prepared alkylchloroaluminum reagents (4 mL, 0.67 M, 2.7 mmol) were added to 4,4’-(thiazole-2,4-diyl)dibenzonitrile (78 mg, 0.27 mmol) in anhydrous toluene (1 mL) and heated to 80 °C overnight under nitrogen. On cooling, the aluminium complex was decomposed by pouring into a slurry of silica gel (2 g) in CHCl_3_. The mixture was stirred for 5 min before filtering, the filter cake was washed with MeOH (20 mL). Removal of solvent gave the crude amidine as a pale solid in quantitative yield. The crude product (100 mg) was purified by reverse phase HPLC using a YMC-pack ODS-A column (250 x 20 mm I.D, 5 µM) eluting with CH_3_CN: Water 0.1% TFA (20-80 % gradient over 20 min). Removal of solvent afforded the desired compound as an off-white solid (38 mg, 22%). Mp 271-272 °C; ^1^H NMR (DMSO, 400MHz) 9.32 (bs, 6H, NH & NH_2_), 8.54 (s, 1H, CH), 8.25 (d, 2H, *J* = 8.5 Hz, ArH), 8.22 (d, 2H, *J* = 8.5 Hz, ArH), 7.92 (d, 2H, *J* = 8.5 Hz, ArH), 7.90 (d, 2H, *J* = 8.5 Hz, ArH); ^13^C NMR (DMSO, 100MHz) 165.5, 165.4, 154.4, 138.7, 137.4, 129.6, 129.2, 127.9, 126.9, 126.8, 119.6; *m/z* (ES) 322 (88 % [M+H]^+^); Found (ES) 322.1121 C_17_H_16_N_5_S requires 322.1126; anal. Found C 46.05, H 3.10, N 12.44, S 5.77, C_21_H_18_N_5_O_4_F_6_S requires C 45.91, H 3.12, N 12.74, S 5.83.

## Chemistry of new compounds from David Boykin laboratory

### General procedure for conversion of nitriles into amidine hydrochlorides (Method A)

To a cold and stirred suspension of the nitrile or dinitrile (0.001 mol) in 15 ml dry THF was added 6.0 ml, (0.006 mol) LiN(TMS)_2_ (1 M in THF), stirred for 24 h, cooled, acidified carefully with saturated ethanolic-HCl, the precipitated white solid stirred for 2 h, solvent removed under reduced pressure, diluted with ether, filtered. The collected solid was added to 10 ml ice water, basified with 2M NaOH, the precipitate was filtered, washed with water and air dried. The solid was suspended in anhydrous ethanol (15 ml) and 5 ml saturated ethanolic-HCl and stirred for 6 h, ethanol was distilled off, triturated with dry ether and filtered. The solid was dried under reduced pressure at 80°C for 12 h to yield (70-75%) amidine hydrochloride.

### General procedure for conversion of nitriles into amidine hydrochlorides (Method B)

A suspension of the nitrile or dinitile (0.001 mole) in 20 ml saturated ethanol-HCl was stirred for 4 days in a closed stoppered flask, followed by precipitation with anhydrous ether. The precipitated light yellow solid imidate ester dihydrochloride was filtered and dried under reduced pressure for 3 h to yield (65% - 70%) amidine hydrochloride. The imidate ester dihydrochloride (0.0005 mole) in 20 ml anhydrous ethanol was saturated with ammonia(g) or 0.03 equivalents of ethylene diamine stirred in ethanol (at reflux) for 12 h, solvent removed, 20 ml ice water added, basified to pH 10 with aqueous 2N NaOH, filtered, and washed with water. The precipitated solid was dried in air, suspended in 10 ml of saturated ethanolic-HCl, and stirred for 2 h. The solvent removed, dry ether 20 ml added, filtered, washed with ether and dried under reduced pressure for 12 h. The product amidine hydrochloride was obtained as yellow solid 60-66% yield.

**4-(5-(4-methoxyphenyl) furan-2-yl) benzimidamide hydrochloride (DB 607)**

A mixture of 5-(4-cyanophenyl)-2-bromo furan^1^ (1.23 g, 0.005 mole) and 4-methoxyphenyl boronic acid (0.93 g, 0.006 mole) in 75 ml dioxane under nitrogen was added K_2_CO_3_ (1.38 g, 0.01 mole, in 5 ml H_2_O), followed by Pd(PPh_3_)_4_ 0.12 g (0.0001 mole) and the solution was heated under reflux for 12-24 h (tlc monitored). The solvent was removed under reduced pressure, solid filtered, washed with hexane and dried in air. The solid was suspended in DCM (100 ml), filtered through celite, concentrated under reduced pressure, triturated with ether: hexane (2:1), and filtered to yield 4-(5-(4-methoxyphenyl)furan-2-yl)benzonitrile as a yellow brown solid 0.76 g (74%) mp 250-2 °C dec; ^1^H NMR (DMSO-d_6_): 7.94 (d, 2H, J= 10.4 Hz), 7.85 (d, 2H, J= 10.4 Hz), 7.78 (d, 2H, J= 10.8 Hz), 7.28 (d, 1H, J= 4.4 Hz), 7.02 (d, 2H, J= 10.8 Hz), 6.97 (d, 1H, J= 4.4 Hz), 3.81 (s, 3H); ^13^C NMR (DMSO-d_6_): 159.3, 154.5, 150.0, 134.1, 132.9, 125.4, 123.5, 122.5, 119.0, 114.4, 111.8, 108.8, 106.9, 55.2; MS: HRMS-ESI-POS: Calcd. for C_18_H_14_NO_2_ *m/z* 276.1024 (M^+^+1), found *m/z* 276.1021. The amidine hydrochloride was obtained as yellow solid (Method A) 0.24 g (74%) ; mp >318°C dec ; ^1^H NMR (DMSO-d_6_): 9.41 (brs, 2H), 9.1 (brs, 2H), 8.01 (d, 2H, J= 8.4 Hz), 7.92 (d, 2H, J= 8.4 Hz), 7.82 (d, 2H, J= 8.4 Hz), 7.324 (d, 1H, J= 3.6 Hz), 7.04 (d, 2H), 7.02 (d, 1H, J= 3.6 Hz), 3.82(s, #H); ^13^C NMR (DMSO-d_6_): 164.9, 159.2, 154.3, 150.2, 134.8, 128.8, 125.5, 125.4, 123.0, 122.5, 114.4, 111.4, 106.8, 55.2; ; MS: HRMS-ESI-POS.: Calcd. for C_18_H_17_N_2_O_2_ *m/z* 293.1289 (M^+^+1), found *m/z* 293.1274; Anal. calcd. for C_18_H_16_N_2_O_2_-HCl: C, 65.75; H, 5.21; N, 8.52; Found: C, 65.78; H, 5.23; N, 8.44.

**4,4’-(thiophene-2,4-diyl) dibenzimidamide dihydrochloride (DB 1077)**

A mixture of 2, 4-dibromothiophene 1.21 g (0.005 mole), 4-cyanophenylboronic acid 1.75 g (0.012 mole) following procedure for DB 607, yielded 2,4-(4-cyanophenyl)thiophene as a yellow solid, 0.86 g (72%) mp >220°C dec ; ^1^H NMR (CDCl_3_): 7.75-7.21 (m, 8H), 7.71 (d, 1H, J= 1.2 Hz), 7.64 (d, 1H, J= 1.2 Hz); ^13^C NMR (CDCl_3_): 143.6, 141.6, 139.4, 138.0, 132.9, 132.8, 126.8, 126.1, 123.8, 123.7, 118.8, 118.6, 111.3, 111.1; MS: HRMS-ESI-POS: Calcd. for C_14_H_11_N_2_S *m/z* 239.0642 (M^+^+1), found *m/z* 239.0639. The diamidine hydrochloride was obtained as yellow solid (Method A) 0.32 g (74%) mp>300°C dec ; ^1^H NMR (DMSO-d_6_): 9.55 (brs, 2H), 9.54 (brs, 2H), 9.31 (brs, 4H), 8.42 (s, 1H), 8.32 (s, 1H), 8.09 (d, 2H, J= 8.4 Hz), 8.03-7.97 (m, 6H); ^13^C NMR (DMSO-d_6_): 164.9, 164.8, 142.5, 140.9, 139.5, 138.3, 129.1, 128.8, 126.6, 126.3, 126.2, 125.4, 125.0, 124.8; MS: HRMS-ESI-POS.: Calcd. for C_18_H_18_N_4_S *m/z* 161.0626 (M^+^+2)/2, found *m/z* 161.0621; Anal. calcd. for C_18_H_16_N_2_S-2HCl-2H_2_O: C, 50.35; H, 5.16; N, 13.05; Found: C, 50.52; H, 5.23; N, 13.22.

**3,3’-(furan-2,5-diyl bis (4,1-phenylene)) dipropanimidamide dihydrochloride (DB 1061)**

To a mixture of 4-bromophenyl propionitrile 0.63 g (0.003 mole) and 2, 5-bis (tributylstannyl) furan in 30 ml anhydrous dioxane under nitrogen was added Pd(PPh_3_)_4_ 0.14 g (0.00012 mole) and the solution was heated under reflux for 12 h (tlc monitored). The solvent was removed under reduced pressure, the solid was filtered, washed with hexane and dried in air. The solid was suspended in DCM (50 ml), stirred 2 h with 20 ml 10% KF (aqueous), the organic layer separated, filtered through celite dried over anhydrous MgSO_4_, filtered, concentrated, triturated with hexane and the solid filtered was filtered to yield 0.34 g (70%) of 3,3’-(furan-2,5-diylbis(4,1-phenylene))dipropanenitrile as a yellow solid mp 120-2°C dec; ^1^H NMR (CDCl_3_): 7.73 (d, 4H, J= 8.4 Hz), 7.30 (d, 4H, J= 8.4 Hz),7.28 (s, 2H), 3.0 (t, 4H, J= 7.6 Hz), 2.66 (t, 4H, J= 7.6 Hz); ^13^C NMR (CDCl_3_): 153.2, 137.2, 130.0, 128.9, 124.4, 119.2, 107.2, 31.5, 19.5; MS: HRMS-ESI-POS: Calcd. for C_22_H_18_N_2_ONa *m/z* 349.1317 (M^+^+Na), found *m/z* 349.1332.

The diamidine dihydrochloride was obtained using Method B: 0.14 g (60%) mp>300°C dec ; ^1^H NMR (DMSO-d_6_): 9.13 (brs, 4H), 8.74 (brs, 4H), 7.64 (d, 4H, J= 8.4 Hz), 7.49 (s, 2H), 7.33 (d, 4H, J=8.4 Hz), 3.0 (t, 4H, J= 7.2 Hz), 2.74 (d, 4H, J= 7.2 Hz); ^13^C NMR (DMSO-d_6_): 170.6, 142.7, 139.4, 132.4, 129.6, 125.8, 125.2, 33.7, 32.03; MS: HRMS-ESI-POS: Calcd. for C_22_H_26_N_4_O *m/z* 181.1053 (M^+^+2)/2, found *m/z* 181.1048; Anal. calc. for C_22_H_24_N_4_O-2HCl-2H_2_O: C, 56.29; H, 6.44; N, 11.93; Found: C, 56.35; H, 6.54; N, 11.86.

**2,5-bis(4-(2-(4,5-dihydro-1H-imidazol-2-yl) ethyl) phenyl) furan dihydrochloride (DB 1062)**

Similarly, 0.245 g (0.005 mole) of the above imidate ester in 20 ml of anhydrous ethanol was allowed to react under reflux (12 h) with 0.06 g (0.0015 mole) ethylene diamine. The solvent was removed under reduced pressure, diluted with water, solid was filtered, dried and converted to dihydrochloride using ethanolic-HCl to yield a yellow solid, 0.15 (62%), mp >325 °C, H NMR (DMSO-d_6_): 8.29 (br, 4H), 8.74 (brs, 4H), 7.77 (d, 4H, J= 8.4 Hz), 7.34 (s, 2H), 7.05 (d, 4H, J=8.4 Hz), 3.0 (t, 4H, J= 7.2 Hz), 2.74 (d, 4H, J= 7.2 Hz); ^13^C NMR (DMSO-d_6_): 170.2, 152.4, 138.5, 128.8, 128.6, 123.6, 108.0, 44.0, 30.5, 27.4; MS: HRMS-ESI-POS: Calcd. for C_26_H_30_N_4_O *m/z* 207.1209 (M^+^+2)/2, found *m/z* 207.1203; Anal. calc. for C_26_H_28_N_4_O-2HCl-2.75H_2_O: C, 58.37; H, 6.68; N, 10.47; Found: C, 58.45; H, 6.54; N, 10.63.

**3,3’-(thiophene-2,5-diylbis(4,1-phenylene)) dipropanimidamide dihydrochloride (DB 1063)**

The dinitrile, 3,3’-(thiophene-2,5-diylbis(4,1-phenylene))dipropanenitrile, was prepared as described for DB1061 yielding a yellow solid 0.77 g (75%); mp 124-6°C dec.;^1^H NMR (CDCl_3_): 7.62 (d, 4H, J= 8.0 Hz), 7.29 (d, 4H, J= 8.0 Hz),7.28 (s, 2H), 3.0 (t, 4H, J= 7.2 Hz), 2.66 (t, 4H, J= 7.2 Hz); ^13^C NMR (CDCl_3_): 143.3, 137.5, 133.5, 129.1, 126.2, 124.2, 119.2, 31.4, 19.4; MS: HRMS-ESI-POS: Calc. for C_22_H_18_N_2_SNa *m/z* 365.1088 (M^+^+Na), found *m/z* 365.1089.

Similarly following the DB 1061 procedure the diamidine dihydrochloride was obtained as a yellow solid 0.16 g (66%), mp >280°C dec; ^1^H NMR (DMSO-d_6_): 9.19 (brs, 4H), 8.77 (brs, 4H), 7.64 (d, 4H, J= 8.0 Hz), 7.51 (s, 2H), 7.33 (d, 4H, J=804 Hz), 2.99 (t, 4H, J= 8.4 Hz), 2.73 (d, 4H, J= 8.4 Hz); ^13^C NMR (DMSO-d_6_):170.0, 142.2, 138.9, 131.9, 129.1, 125.3, 124.7, 33.2, 31.6; MS: HRMS-ESI-POS: Calcd. for C_22_H_26_N_4_S *m/z* 189.0939 (M^+^+2)/2, found *m/z* 189.0931; Anal. calcd. for C_22_H_24_N_4_S-2HCl-1.5H_2_O: C, 55.45; H, 6.13; N, 11.76; Found: C, 55.52; H, 6.34; N, 11.63.

**2, 5-bis(4-(2-(4,5-dihydro-1H-imidazol-2-yl) ethyl) phenyl) thiophene dihydrochloride (DB 1064)**

Similarly following the procedure for DB1062 the diamidine dihydrochloride was obtained as a yellow solid, 0.17 g (62%); mp >225°C dec.; ^1^H NMR (DMSO-d_6_): 10.19 (s, 4H), 7.63 (d, 4H, J=7.6 Hz), 7.5 (s, 2H), 7.30 (d, 4H, J= 7.6 Hz), 3.78 (s, 8H),2.97 (t, 4H, J= 6.4 Hz), 2.81(t, 4H, J= 6.4 Hz); ^13^C NMR (DMSO-d6): 170.2, 140.2, 138.9, 131.9, 129.1, 125.4, 124.8, 44.1, 30.6, 27.4; MS: HRMS-ESI-POS: Calcd. for C_26_H_29_N_4_S *m/z* 429.2113 (M^+^+1), found *m/z* 429.2109; Anal. calcd. for C_26_H_28_N_4_S-2HCl-3.0H_2_O: C, 56.21; H, 6.53; N, 10.08; Found: C, 56.34; H, 6.61; N, 10.24.

**4-(5-(1-methyl-1H-benzo[d]imidazol-2-yl) furan-2-yl) benzimidamide dihydrochloride (DB 960)**

To a stirred solution of 5-(4-cyanophenyl) furan-2-aldehyde^2^ 1.23 g (0.005 mole), 1-amino-2-*N*-(methylamino) benzene 0.61 g (0.005 mol) in 20 ml dry DMF under N_2_ was added sodium metabisulfite 0.95 (0.005 mol) and the mixture was heated at 130°C for 12 h (tlc monitored). The solvent was removed, the residue was triturated with cold water, separated solid was filtered, washed with water and air dried. The solid was stirred with 1:1 mixture of DCM-ether, filtered and dried in vac at 70°C for 4 h to give the nitrile, 4-(5-(1-methyl-1H-benzo[d]imidazol-2-yl)furan-2-yl)benzonitrile, as a yellow brown solid, 1.1 g (72%), mp >290°C dec ; ^1^H NMR (DMSO-d_6_): 8.03 (d, 2H, J= 8.4 Hz), 7.92 (d, 2H, J= 8.4 Hz), 7.69-7.63 (m, 2H), 7.46 (d, 1H, J= 3.6 Hz), 7.40 (d, 1H, J= 3.6 Hz), 7.34-7.23 (m, 2H), 4.13 (s, 3H); ^13^C NMR (DMSO-d_6_): 152.4, 145.9, 143.2, 142.4, 136.0, 133.2, 132.9, 124.2, 122.7, 122.2, 118.9, 118.5, 114.7, 111.1, 110.2, 109.9, 31.4; MS: HRMS-ESI-POS: Calcd. for C_19_H_14_N_3_O *m/z* 300.1136 (M^+^+1), found *m/z* 300.1132.

The amidine hydrochloride was obtained as yellow solid (Method B) 0.3 g (78%); mp >300°C dec; ^1^H NMR (DMSO-d_6_): 8.09 (d, 2H, J= 8.7 Hz), 7.90 (d, 2H, J= 8.7 Hz), 7.78-7.70 (m, 2H), 7.61 (d, 1H, J= 3.9 Hz), 7.48-7.42 (m, 2H), 7.45 (d, 1H, J= 3.9 Hz), 4.13 (s, 1H); ^13^C NMR (DMSO-d_6_): 165.5, 155.6, 141.5, 141.3, 135.3, 134.8, 133.9, 129.5, 128.0, 126.0, 125.8, 125.5, 119.6, 116.4, 112.4, 112.0, 32.9; MS: HRMS-ESI-POS: Calcd. for C_19_H_18_N_4_O *m/z* 159.0740 (M^+^+2)/2, found *m/z* 159.0733; Anal. calcd. for C_19_H_16_N_4_O-2HCl-1H_2_O: C, 56.02; H, 4.94; N, 13.76; Found: C, 56.18; H, 4.91; N, 13.51.

